# Cross-species blood transcriptional correlates of BCG-mediated protection against tuberculosis include innate and adaptive immune processes

**DOI:** 10.1101/2025.05.05.652268

**Authors:** Kate Bridges, Denis Awany, Anele Gela, Temwa-Dango Mwanbene, Christopher M Sassetti, Thomas J Scriba, Douglas A Lauffenburger

## Abstract

The immune mechanisms induced by the Bacillus Calmette-Guérin (BCG) vaccine, and the subset of which mediate protection against tuberculosis (TB), remain poorly understood. This is further complicated by difficulties to verify vaccine-induced protection in humans. Although research in animal models, namely mice and non-human primates (NHPs), has begun to close this knowledge gap, discrepancies in the relative importance of biological pathways across species limit the utility of animal model-derived biological insights in humans. To address these challenges, we applied a systems modeling framework, Translatable Components Regression (TransCompR), to identify human blood transcriptional variability which could predict *Mtb* challenge outcomes in BCG-vaccinated NHPs. These protection-associated pathways included both innate and adaptive immune activation mechanisms, along with signaling via type I interferons and anti-mycobacterial T helper cytokines. We further partially validated the associations between these mechanisms and protection in humans using publicly available microarray data collected from BCG-vaccinated infants who either developed TB or remained healthy during two years of follow-up. Overall, our work demonstrates how species translation modeling can leverage animal studies to generate hypotheses about the mechanisms that underlie human infectious disease and vaccination outcomes, which may be difficult or impossible to ascertain using human data alone.

## INTRODUCTION

Tuberculosis (TB) is presently the deadliest infectious disease caused by a single agent, with 10.8 million incident cases and 1.25 million deaths globally in 2023 (1). Among preventive interventions, the only TB vaccine available is the Bacillus Calmette-Guérin (BCG) vaccine (2), which is typically administered intradermally at birth in high incidence countries. Despite efficacy against disseminated disease in young children, BCG confers variable protection against adolescent and adult pulmonary TB (3–5) via mechanisms that remain poorly understood by the field. Robust vaccine-mediated immune correlates of protection need to be identified to better inform the development and use of TB vaccines.

A major challenge with TB vaccine research is the difficulty in assessing vaccine-mediated protection in humans. At minimum, studying post-vaccination clinical outcomes in humans requires clinical follow-up of thousands of participants (6–8) where exposure to pathogen is not always a known variable. Moreover, diagnostic tools for TB vary widely in their sensitivity, specificity, cost, portability, and resource intensity (9–11), which complicate their application to large human cohorts in different ways and limit disease classification. Some clinical outcomes, such as bacterial burden in lung tissues, are simply not directly observable in living humans given present diagnostic tools and ethical consideration (12–14).

For these reasons, TB vaccine research heavily relies on the use of animal models, including mice and non-human primates (NHPs), to evaluate vaccine-mediated immune mechanisms (15,16). However, animal model-derived biological insights are not guaranteed to be directly translatable to humans (17,18), owing in part to divergent immune processes (19,20) and to discrepancies in the phenotypic relevance of conserved immune processes (21) across species. For example, the long-held paradigm regarding the mechanism of action of BCG was that protective immunity is conferred by vaccine-induced IFNγ-producing CD4^+^ T cells (22), which was supported in part by mouse studies (23,24). However, various clinical studies have asserted little to no correlation between IFNγ^+^CD4^+^ T cells and BCG-mediated protection (6,25), suggesting that cross-species differences could be one factor which limits the translational potential of TB vaccine studies in animal models. This example in particular highlights the need for the development and widespread use of improved approaches to enhancing translation proficiency (26–28).

Our group previously developed a computational framework called Translatable Components Regression (TransCompR), which was designed to relate orthogonal axes of transcriptional or proteomic variability in one species to disease biology and measurable phenotypes observed in another species (27). In doing so, TransCompR explicitly accounts for differences in the relative importance of disease-relevant processes and pathways in the two species. Although originally applied in the contexts of inflammatory bowel disease (27,29) and neuropathology (30,31), we have recently used this framework to identify transcriptional variability in mouse TB models which could best predict human TB phenotypes (21,32), demonstrating the utility of TransCompR to uncover species-translatable signatures in an infectious disease context.

In this work, we adapted TransCompR to identify human-relevant biological pathways which are associated with the outcome of post-*Mtb* challenge in BCG-vaccinated NHPs. We show that a linear mathematical model built on human blood transcriptomic data can predict vaccine-mediated protection against TB as assessed in publicly available data from NHP cohorts (33,34). These protection-associated directions of human transcriptional variance corresponded to the activation of innate and adaptive immune mechanisms, including type I IFN and anti-mycobacterial T helper cytokine signaling. We further partially validated the associations between these directions of transcriptional variance and protection using publicly available microarray data collected from a heterogeneous cohort of BCG-vaccinated infants who either developed TB or remained healthy during two years of follow-up (35). Overall, we demonstrate how translational cross-species modeling can leverage animal studies to uncover mechanisms of action in humans where no outcome data are available.

## RESULTS

### Bulk blood transcriptomics data collection to characterize BCG-induced biological variability across species

To develop an understanding of vaccine-induced immune responses in animal models and humans, we used non-human primate (NHP) and human bulk blood transcriptomic data. We included data from 60 South African infants who received percutaneous or intradermal BCG vaccination at birth and were selected from a parent cohort study of 11,680 infants (6,7) to reflect a range of BCG responses based on antigen-induced T cell IFNγ production (see **Methods**). Bulk transcriptional profiling on peripheral blood mononuclear cells (PBMCs) and intracellular cytokine staining (ICS) on whole blood collected at 10 weeks of age were performed for characterization of peripheral immune responses (**Figs. 1A and S1A**). The NHP dataset was previously published in a study by Liu *et al* (33). NHPs in this cohort [n = 34; ‘dose cohort’ (36)] were vaccinated intravenously with BCG across a range of doses, and whole blood was collected for transcriptional profiling at baseline and at four post-vaccination timepoints (**Fig. 1B**). Plasma and bronchoalveolar lavage (BAL) samples were also collected from these NHPs at these or similar timepoints for flow cytometry and antibody profiling (37). At 24 weeks post-vaccination, these NHPs were challenged with *Mtb*, and outcome was assessed at 8-12 weeks post-*Mtb* challenge via quantification of bacterial burden (as colony forming units, or CFU) in lung tissue.

**Figure 1.**
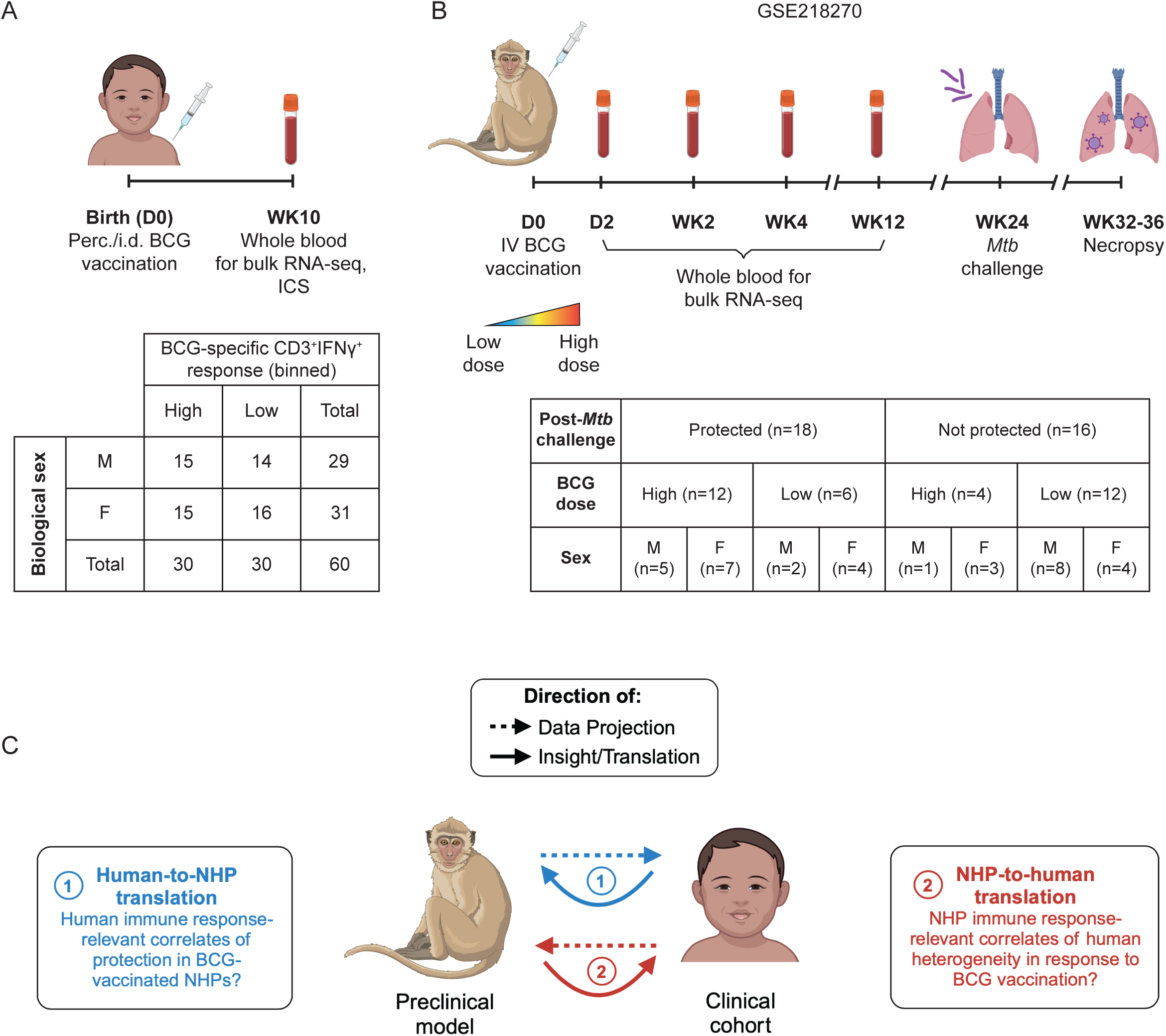
BCG-induced patterns in gene expression across species. (A) (Top) Schematic describing collection of blood for bulk RNA-seq and profiling with intracellular cytokine staining (ICS) in a cohort of 60 BCG-vaccinated South African infants. (Bottom) Table detailing breakdown of 60 human samples by biological sex and binned BCG-specific CD3^+^IFNγ^+^ response as assessed by ICS. (B) (Top) Schematic describing collection of blood for bulk RNA-seq in a cohort of 34 IV BCG-vaccinated non-human primates (NHPs). Vaccination was administered over a range of BCG doses. The bulk RNA-seq data was previously published at GSE218270. (Bottom) Table detailing breakdown of 34 NHP samples by biological sex, administered binned BCG dose, and binned post-*Mtb* challenge lung bacterial burden (‘protected’ = log_10_(CFU) < 2; ‘not protected’ = log_10_(CFU) > 2). (C) Schematic conceptually describing species translation. Translation modeling involving two species presents two potential directions for analysis, which affects the biological questions that can be probed computationally.

### TransCompR model identifies human transcriptional variability associated with BCG-mediated protection in non-human primates

To construct a species translation model, we used Translatable Components Regression [TransCompR (27)], a method previously developed in our group to relate axes of data modality-derived variance in one species to a phenotype of interest in another species. Like other species translation approaches, TransCompR analyses have directionality (**Fig. 1C**). Historically, TransCompR has been used to relate transcriptional variability in mouse or NHP models of disease to human disease characteristics [e.g., animal model-to-human translation (21,27,38); visualized in **Fig. 1C, right**, in red] with the objective being to better understand axes of variance in animal models with respect to their human relevance. However, human-to-animal model translation (visualized in **Fig. 1C, left**, in blue), in which human variance is regressed against biological outcomes as assessed in animal models, also offers actionable insights. This direction of species translation is particularly useful for studies in which assessing individuals’ responses to immunomodulation is difficult or limited by experimental or ethical bounds, a point that our group has previously demonstrated in non-infectious disease contexts (29,30). Different directions of translation address different scientific questions; therefore, depending on the application, one of the two possible directions might address a more informative question than the other.

Because vaccine-induced protection is challenging to ascertain in humans, in this work we chose to pursue human-to-NHP translation modeling (**Fig. 1C, left**, *in blue only*) to investigate human immune response-relevant correlates of protection in BCG-vaccinated NHPs. To do this, we first identified 9,628 gene homologs between the human and NHP datasets for inclusion in our cross-species model (**Fig. 2A**). We then used principal component analysis (PCA) on the human bulk RNA-seq data to construct a latent space which describes orthogonal directions of transcriptional variance in the South African infant cohort. The first 15 human-derived principal components (hPCs), which captured approximately 80% of total variance in the infant cohort (**Fig. 2B**), were retained for cross-species modeling.

**Figure 2.**
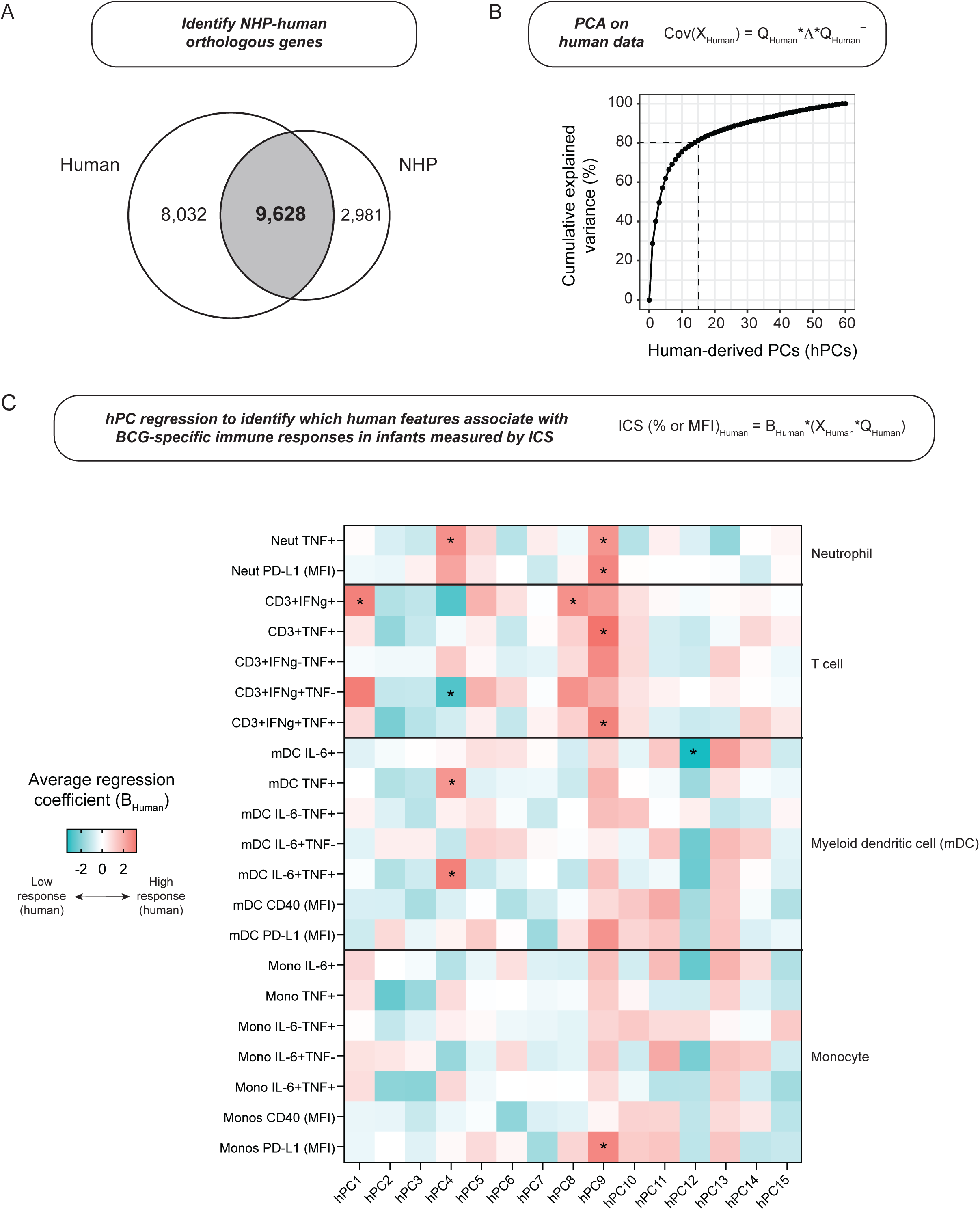
Human-derived principal components associate with BCG-specific immune responses in infants. (A) Venn diagram demonstrating the number of genes, as profiled in the human and NHP bulk RNA-seq datasets, that were either species-specific or orthologous. (B) Line plot demonstrating cumulative explained variance (%) across subsequent human-derived principal components (hPCs) as calculated on the human bulk RNA-seq dataset which was first limited to only orthologous genes. (C) Heatmap of univariate regression coefficients from individual human-derived principal components regressed against patterns in cytokine expression (% unless otherwise indicated) from immune cells stimulated with BCG *ex vivo*. Regression coefficients are averaged across results from 5-fold cross validation. *p < 0.05 across folds. For each fold, p-values are calculated by Wald test.

Prior to translation, we were first interested to understand how these hPCs, representative of blood transcriptional variance, related to measurable human phenotypes. To do this, we univariately regressed each of the 15 hPCs retained in our model against patterns in cytokine expression by T cells, monocytes, myeloid dendritic cells (mDCs), or neutrophils, assessed by ICS on whole blood and collected at 10 weeks post-BCG priming, that was either left unstimulated or stimulated *ex vivo* with BCG or staphylococcal enterotoxin B (SEB), a positive control (**Figs. 2C and S1A-B**). Associations quantified by regression were notably different between SEB– and BCG-stimulated cells. We further found 5 of these 15 hPCs to be significantly associated with one or more BCG-specific cytokine-expressing cellular subsets. hPC1, hPC4, and hPC8 were significantly associated with BCG-specific IFNγ-expressing T cells, while hPC9 was instead significantly associated with BCG-specific TNF-expressing T cells.

Interestingly, hPC12 was associated with IL-6-expressing mDCs, suggestive of human transcriptional variance which might correlate with innate responses to BCG. Because antigen-specific IFNγ production by T cells is typically considered a protective immune marker in the context of BCG vaccination (22), we binned infants into high and low responder groups by their BCG-specific IFNγ^+^ T cell response (**Fig. S2A**) and used univariate logistic regression to identify hPCs predictive of responder group. Separation between responder groups was not observed along the first two hPCs (**Fig. S2B**).

Instead, hPC4, hPC8, and hPC9, which collectively explained 12% of total transcriptional variance, were found to significantly distinguish infants by responder group (**Fig. S2C-D**). Using covariate logistic regression, these 3 hPCs were capable of predicting responder group with high classification accuracy (**Fig. S2E**), signaling that, perhaps surprisingly, hPCs which explain *less* transcriptional variance overall might actually be more relevant to human immunological responses of interest.

To next approach species translation, we then projected each NHP sample collected at each post-vaccination timepoint into our hPC space using the gene ortholog loadings for each of the 15 hPCs retained in our model, resulting in NHP sample scores along these hPC axes. This projection allowed us to ‘reorder’ or ‘filter’ the NHP data to reflect the most relevant directions of variance in the human data. As expected, for human samples, the variance explained by each subsequent hPC monotonically decreased, while this trend was broken for projected NHP samples (**Fig. 3A, left**).

**Figure 3.**
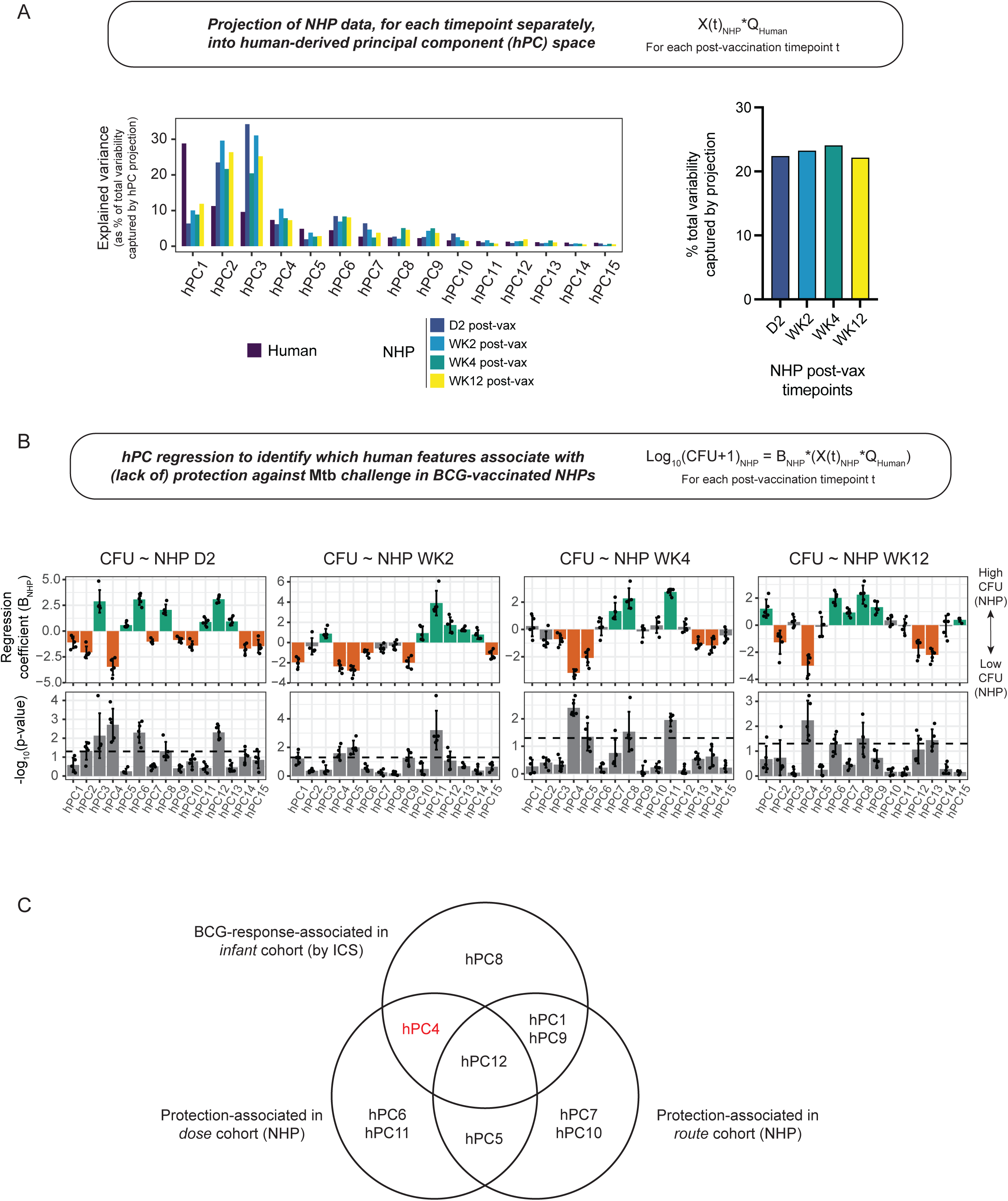
Human-derived principal components can also predict post-*Mtb* challenge bacterial burden in NHPs. (A) Bar graphs showing explained variance captured by projecting the NHP bulk RNA-seq data, restricted to orthologous genes, into hPC space (right) as a percentage of total variance preserved for each post-vaccination NHP dataset and (left) as distributed across each hPC in the TransCompR model. (B) Bar graphs showing (top) 5-fold logistic regression coefficients (summary data presented as mean ± SD) and (bottom) distribution of corresponding p-values from univariate regression of each hPC against post-*Mtb* challenge bacterial burden (CFU). For each fold, p-values are calculated by Wald test. The dotted line denotes p = 0.05. (C) Venn diagram detailing overlap of hPCs which were found to be significantly associated with protection in the dose and route cohorts of NHPs and with BCG-specific immune cytokine expression in the South African infant cohort. hPC4 is highlighted in red owing to its unexpected correspondence between high bacterial burden in NHPs and high BCG-specific T cell response in infants.

Interestingly, while patterns in explained variability were similar for each NHP timepoint, they are not identical, which hinted at how axes of variance in the data might be changing over time. Moreover, across post-vaccination timepoints in the NHP cohort, the hPC projection preserved approximately 25% of the total variance in the NHP datasets (**Fig. 3A, right**) which reemphasized the idea that common biological pathways and processes may differentially contribute to transcriptional variability, and, ultimately, higher order biological function, across species.

Because many immune responses that were initially quantified in the dose cohort of NHPs exhibited dose dependence (4), we were first interested to understand how dose-linked transcriptional variance might be reflected in the hPC-projected bulk RNA-seq data. To do this, we univariately regressed the hPC scores from the projected NHP samples against the dose of vaccination administered to each NHP. Many hPCs were significantly associated with BCG dose (**Fig. S3**), both with and without time dependence, suggesting that dose-linked variation was reflected at the transcriptional level and that this variation was preserved upon hPC projection. This preservation also raised the possibility that the range of immune responses generated in NHPs by varying BCG dose maintained some human relevance. hPC4 was also significantly associated with BCG dose in NHPs at all post-vaccination timepoints, suggesting that this component captures variance related to robust and sustained response mechanisms that are potentially core and conserved in both species.

Finally, toward our goal of identifying vaccine-induced correlates of protection in the NHP data which are human-relevant, we then, for each post-vaccination timepoint separately, univariately regressed the hPC scores from the projected NHP samples against their post-*Mtb* challenge bacterial burden to test for statistically significant, time-dependent association (**Fig. 3B**). This analysis identified 5 protection-associated hPCs (with time dependence): hPC4 (all post-vaccination timepoints); hPC5 (2wk); hPC6 (2d); hPC11 (2wk and 4wk); and hPC12 (2d). These associations largely held whether post-*Mtb* challenge outcome was treated as a continuous (CFU) or discrete (‘protected’ versus ‘not protected’) variable, and covariate regression using each time-specific subset of protection-associated hPCs confirmed significant improvement in prediction of post-challenge outcome compared to null models with random hPCs or shuffled outcomes (**Fig. S4**).

To examine the robustness of these associations between hPCs in our model and protection against TB, we repeated the projection and regression steps of our species translation approach using an independent cohort of NHPs (**Fig. S5A-B**).

Referred to as the ‘route cohort’ (33,34), these NHPs (n = 34) were vaccinated with BCG via different routes, and, similarly to the dose cohort, blood was collected for transcriptional profiling at baseline and at three post-vaccination timepoints, with later *Mtb* challenge and necropsy to assess protection against disease. Like the dose cohort, hPC projection preserved approximately 20-25% of the total variance in the NHP route cohort datasets (**Fig. S5A**). Regressing hPC-projected route cohort samples against post-*Mtb* challenge bacterial burden provided further support for hPC5 (2wk) and hPC12 (2d) as time-dependent, species-translatable axes of variance which are associated with protection against TB across independent NHP cohorts (**Fig. S5B-C**).

Finally, we examined the overlap between hPCs significantly associated with protection across NHP cohorts and hPCs significantly associated with BCG-specific immune responses in infants (**Fig. 3C**). Perhaps expectedly, three T cell response-associated hPCs (hPC1, hPC4, and hPC9) were also significantly associated with protection in one of the two NHP cohorts, although, contradictorily, for hPC4, protected NHP samples were separated on the same end of the axis as the low T cell responders (**Fig S4A**; compare to **Fig. S2D**). Of the hPCs found to be protection-associated in both NHP cohorts, hPC5 did not correspond to any patterns in human immune cytokine expression that were assessed by ICS, while hPC12 was instead associated with antigen-reactive innate responses in infants. Overall, our cross-species modeling identified axes of transcriptional variability which maintained significant associations with measurable human and NHP biological outcomes.

### Cross-species correlates of protection represent biological axes related to innate, adaptive, and humoral immunity

We next wanted to biologically interpret these species-translatable hPCs as they represent orthogonal directions of variance, and therefore we expected them to reflect distinct (although not wholly disjoint) biological processes. We focused on hPC5 and hPC12, which were protection-associated across both NHP cohorts, for these analyses. To manage this interpretation, we used gene set enrichment analysis (GSEA) to compare the gene loadings for these hPCs against gene sets in the Reactome Pathways and Kyoto Encyclopedia of Genes and Genomes (KEGG) databases (39,40). Using Reactome as the reference database, hPC5 (2wk) was significantly enriched (p_adj_ < 0.1) for 175 terms, while hPC12 (2d) was significantly enriched for only 8 terms (**Fig. 4A, Table S1**). With KEGG, hPC5 and hPC12 were respectively significantly enriched for 35 and 8 immune– and cell function-related terms (**Fig. S6**). Across reference databases, we noted enrichment of hPC12 for innate immune activation, antiviral immunity, and type I IFN signaling, mechanisms which have been shown to be altered by BCG, *Mtb* (41), and other pulmonary infections (42), while hPC5 was enriched for signaling pathways related to innate-adaptive immune crosstalk, T helper differentiation, and B cell activation (**Figs. 4B and S6**). Included in these terms enriched along hPC5 was Dectin-1 signaling. Dectin-1 is a C-type lectin that functions as a pattern recognition receptor, is primarily expressed on antigen-presenting cells [APCs (43)], including dendritic cells (DCs) and B cells, and has been described to play a role in immune responses to mycobacteria, including *Mtb* and BCG (44,45). Signaling downstream of Dectin-1 activation is known to induce production of reactive oxygen species (ROS) in neutrophils, as well as NFκB– and MAPK-mediated expression of proinflammatory cytokines, including TNF, IL-12, IL-6, and IL-1β, in APCs (43,46). Because IL-12 and IL-6/IL-1β are known to direct the differentiation of naïve CD4^+^ T cells into T helper (Th) subsets Th1 and Th17, respectively (47), it is possible that hPC5 represents an axis of transcriptional variability that is associated with Dectin-1-mediated coupling of innate and adaptive immunity.

**Figure 4.**
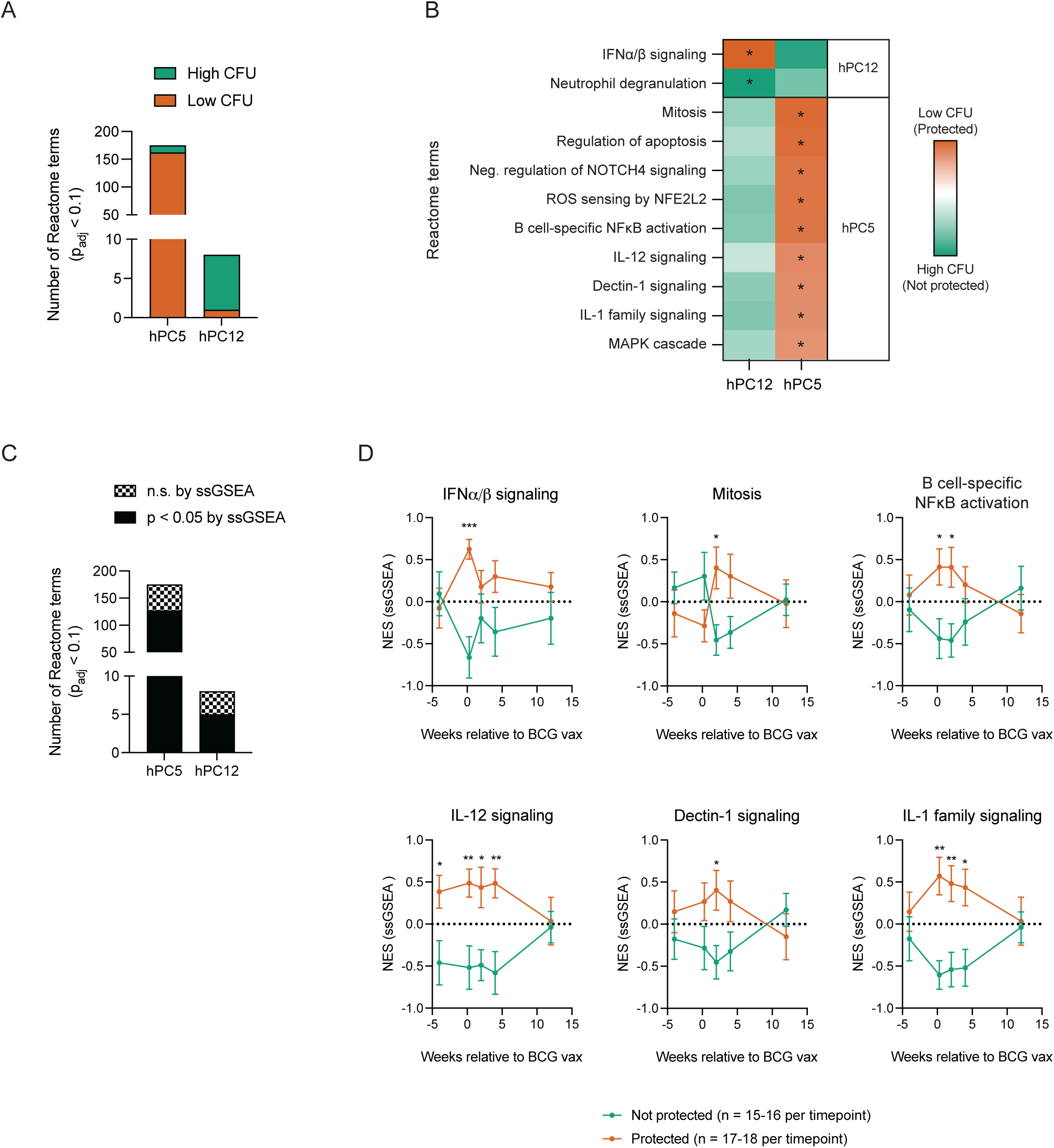
Protection-associated hPCs are enriched for innate and adaptive immune processes and pathways. (A) Bar graph demonstrating the total number of significantly enriched (p_adj_ < 0.1) Reactome terms along hPC5 and hPC12. (B) Heatmap showing enrichment scores for select Reactome terms across hPC5 and hPC12. An asterisk denotes where enrichment scores were deemed statistically significant (p_adj_ < 0.1). P-values were estimated using an adaptive multi-level split Monte-Carlo scheme implemented in the fgsea package in R. (C) Bar graph as in (A), with terms now separated into significant (black) or non-significant (checkered) categories by how their leading-edge gene sets distinguished NHP samples by challenge outcome (p-values obtained by unpaired t-test). hPC12-enriched terms were evaluated for their ability to separate NHP samples collected 2 days post-vaccination by their challenge outcome, while hPC5-enriched terms were evaluated using NHP samples collected 2 weeks post-vaccination. (D) Normalized enrichment scores (NES) calculated using single sample gene set enrichment analysis (ssGSEA) for each sample in the NHP dose cohort. Gene sets for each pathway/process were derived from the Reactome Pathway Database. Sample scores were then lumped by protection outcome as evaluated post-*Mtb* challenge for visualization. Data is presented as mean ± SEM. *p < 0.05, **p < 0.01, ***p < 0.001 by unpaired t-test with Holm-Šídák’s multiple comparison correction as calculated in GraphPad.

To confirm that these hPC-enriched Reactome pathways and processes separated NHPs by their post-*Mtb* challenge infection status, we used single-sample GSEA (ssGSEA) to calculate enrichment scores for these gene sets for each individual sample in the NHP dose cohort at each profiled timepoint and compared between ‘protected’ and ‘non-protected’ NHPs. As expected, we observed statistically significant, time-dependent separation of samples by the majority of hPC12– and hPC5-enriched terms (**Fig. 4C**), including the interferon and cytokine signaling pathways that we highlighted (**Fig. 4D**), at one or more post-vaccination timepoints. Repeating this analysis in the route cohort of NHPs, we observed similar patterns in time-dependent enrichment which distinguished protected from non-protected NHPs (**Fig. S7**). Overall, interpreting protection-associated hPCs with gene set enrichment analyses identified species-translatable biological pathways and processes which were associated with BCG-mediated protection in NHPs.

### Independent cohort of South African infants with two-year follow-up outcomes partially validates species-translatable correlates of protection

Finally, because our cross-species analyses allowed us to generate hypotheses about transcriptional correlates of protection in vaccinated infants, we wanted to approach their validation. Although vaccine-induced protection is difficult to ascertain in humans, Fletcher *et al* (35) previously published a microarray dataset which captured PBMCs, either left unstimulated or stimulated *ex* vivo with BCG, from South African infants who were vaccinated with BCG at birth and who later either developed TB (cases) or who remained disease-free (controls) during two years of follow-up (**Fig. 5A**). While the original study failed to reliably identify a correlate of risk, the authors instead classified patients into two clusters based on patterns in gene expression (**Fig. 5B**), where immune phenotypes presented differently in cases depending on their cluster membership. Therefore, we were interested to use this microarray dataset with case and control outcomes as a validation set. To do this, we calculated enrichment scores for each sample using the top 5% of genes along each extreme of hPC5 and hPC12 and compared these scores across clusters and follow-up outcomes for each *ex vivo* stimulation condition that was profiled (**Fig. 5C and Table S2**). Although no meaningful separations were observed by hPC5 gene expression, we noted, for Cluster 1 only, that cases and controls could be nearly significantly distinguished by expression of genes in the top 5% of (+) hPC12 across most individual *ex vivo* stimulation conditions. Using Fisher’s method to merge *P* values across all stimulation and timepoint pairs (four conditions total) indeed yielded statistical significance for this comparison only (*P* = 0.027; see **Table S3**). This result lent support for the idea that our species translation modeling can leverage animal model data to identify correlates of relevant human phenotypes that can be difficult to assess using human data alone.

**Figure 5.**
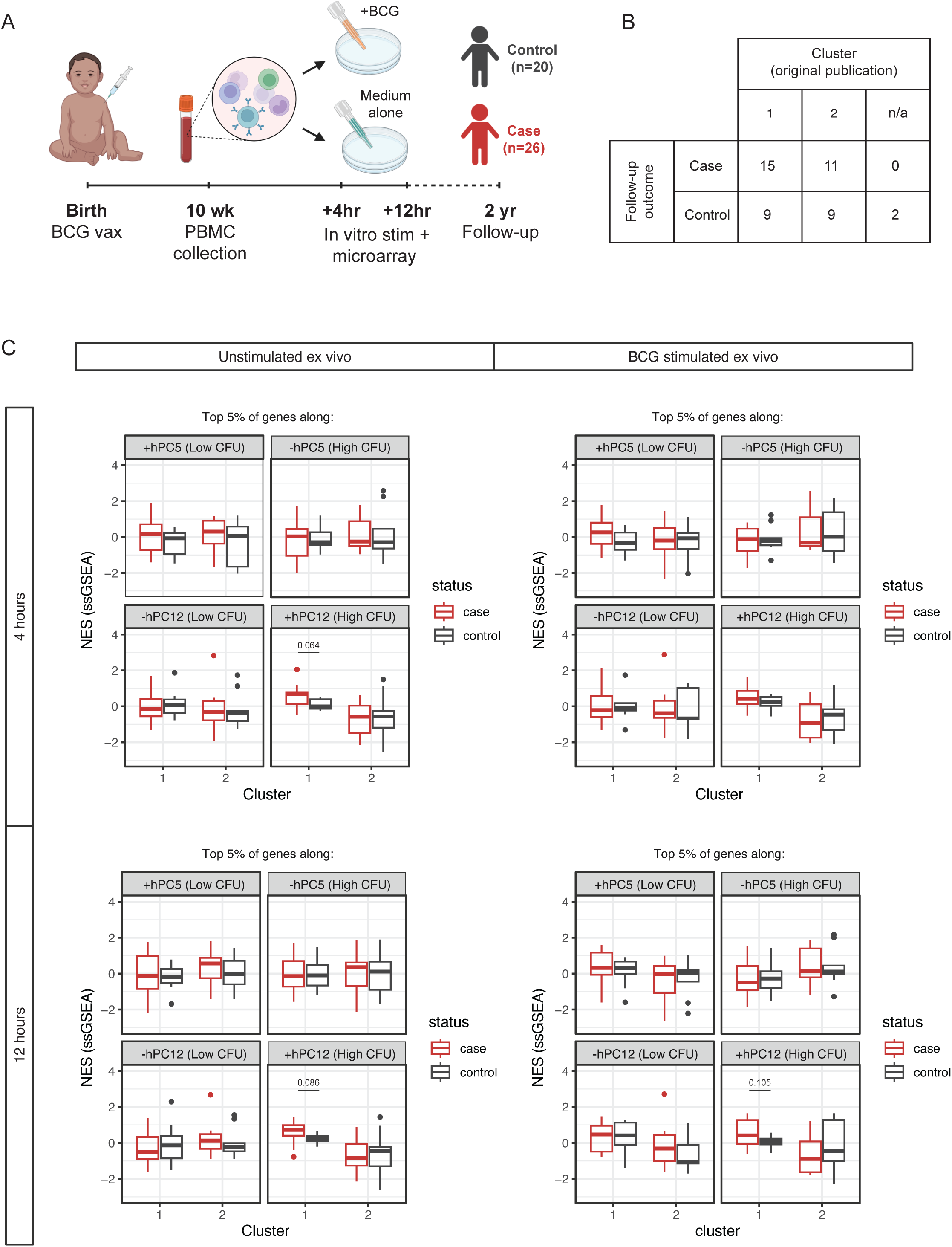
GSE20716 provides partial validation for associations between hPCs and protection. (A) Schematic detailing microarray data collection strategy for cohort of South African infants which is publicly available at GSE20716. (B) Table describing breakdown of 46 patient samples in (A) by two-year follow-up outcome and by assigned cluster from original publication. (C) Boxplots visualizing normalized enrichment scores (NES) calculated with ssGSEA for each microarray sample either (left) left unstimulated or (right) BCG-stimulated *ex vivo* for (top) 4 or (bottom) 12 hours. The thick center line of the box denotes the median; the bounds of the boxes (hinges) denote the 25^th^ and 75^th^ percentiles; the upper and lower whiskers extend ± 1.5 * interquartile range from the upper and lower hinges, respectively; data beyond the whiskers are considered outliers and plotted individually. Samples were separated by case (red) versus control (blue) follow-up outcome and by major cluster assigned in the original publication of this dataset. Statistical testing between distributions of enrichment scores between cases and controls was conducted by unpaired t-test.

## DISCUSSION

BCG is currently the only TB vaccine available. Despite having been first approved for clinical use over a century ago (2), our understanding of the mechanism(s) of action of BCG and the limitations of its efficacy remain elusive (3–5), likely due in part to constraints on disease characterization imposed by present clinical diagnostic tools (10,11,48). Animal models such as mice and NHPs have thus been employed to investigate BCG-mediated protective immunity, but these models can further complicate the identification of human-translatable biological insights due to cross-species discrepancies in the phenotypic relevance of immune processes and pathways (17,18,21).

Our lab has developed a cross-species modeling framework called TransCompR (27) which evaluates orthogonal axes of transcriptional or proteomic variability in one species with respect to a phenotype or biological outcome of interest in another species, allowing for the explicit accounting of cross-species differences. Previous applications of TransCompR have identified underappreciated pathological mechanisms in various disease contexts which were not apparent with traditional cross-species comparisons (21,27,29–32). Here, we adapted TransCompR to identify human-relevant biological pathways which were predictive of BCG-mediated protective immunity, a disease-relevant biological outcome which is best characterized in animal models given experimental and ethical considerations. We used both novel and publicly available (33,34) bulk blood transcriptomics datasets from human participants and animal models to construct and test our species translation model. It is important to note that lung-localized immune responses, collected at the site of infection and typically quantified via BAL, might be more disease-relevant. However, blood-based responses maintain clinical relevance because they are more easily measured at scale in human participants.

Our human-derived principal component model, built on the human blood transcriptomic data, was separately able to predict *ex vivo* BCG-specific immune responses in infants (**Fig. 2**) and post-*Mtb* challenge outcomes as assessed in two different publicly available, BCG-vaccinated NHP cohorts (**Fig. 3**). Interestingly, we found no hPCs which were associated both with protection across the two NHP cohorts and BCG-specific T cell responses in infants. This result is somewhat surprising given that antigen-specific IFNγ production by T cells has been historically considered to be a BCG-mediated immune correlate of protection (22,49), although there is a growing body of literature which contradicts this in both humans and animal models (6,50). It is also possible that BCG-mediated antimycobacterial immunity requires polyfunctional T cell responses, which have been demonstrated by previous literature (51) and which are not captured by observing IFNγ expression alone.

Instead, we observed a significant association between hPC12 and antigen-reactive IL-6 expression in myeloid DCs. IL-6 is a proinflammatory cytokine known to facilitate innate-adaptive immune crosstalk, and it has been shown to drive CD4^+^ (Th1/17) and CD8^+^ T cell activation in the contexts of BCG vaccination and *Mtb* infection (52,53). However, the observed antigen “specificity” of this response, given that hPC12 maintained no significant associations with IL-6^+^ mDCs (%) under unstimulated or SEB-stimulated conditions (**Fig. S1B**), was unexpected given that it is an innate immune response despite its ties to adaptive immunity. This finding has parallels to recent work demonstrating BCG-induced innate immune memory, also called *trained immunity*, primarily via epigenetic reprogramming in circulating monocytes (54–57). This phenomenon has also been observed in mature myeloid cell types, including lung-resident alveolar macrophages with BCG vaccination (58) and splenic DCs with fungal pathogenic stimulation (59). More evidence is needed to definitively disentangle whether DCs can be induced to elicit memory-like immunological responses against TB and how these responses might participate in BCG-mediated protective immunity.

Of those human-derived PCs found to associate with protection in NHPs, a common theme of time dependence emerged. This was a fundamental nuance of our analyses which emphasized the dynamic nature of vaccine-induced processes and pathways and highlighted the importance of considering time as a variable when assessing BCG-mediated immune correlates of protection. On this topic, we further noted that the temporal component of the associations between hPC5 and hPC12 and protection was the same for both cohorts (**Figs. 3B and S5B**), lending further support for the idea that these orthogonal axes of transcriptional variance represent species-translatable, time-dependent, and largely acute mechanisms which play a role in BCG-mediated protective immunity.

We next used gene set enrichment analysis (GSEA) to reveal which biological processes and pathways might be captured by the protection-associated human-derived PCs in our species translation model, focusing our interpretation on hPC12 and hPC5. hPC12, across reference databases (39,40), was enriched for innate immune activation via RIG-I-like and NOD-like receptors, antiviral immunity, and type I IFN signaling. RIG-I agonism in concert with BCG vaccination has been shown, albeit in mice, to enhance antigen presentation, myeloid-specific IFN-β secretion, memory T cell expansion, and BCG-mediated protective responses (60), and therefore it is possible that hPC12 captures transcriptional variability associated with multiple innate immune processes which ultimately cooperate to facilitate anti-TB immunity through the initiation of mycobacterial-specific T cell responses. The characterization of type I IFN signaling as protective might be controversial given that previous studies have described this pathway as an immune correlate of risk (61,62); however, timing and vaccination status might provide crucial context, as other studies have described a protective role for type I IFNs against *Mtb* infection specifically during the acute phase of response to BCG vaccination (33,63).

Regarding hPC5, we found this human-derived axis of transcriptional variability to be enriched for biological processes concerning innate-adaptive immune crosstalk, T helper differentiation, and B cell activation. Of particular interest was the enrichment of hPC5 for signaling via Dectin-1, which is typically restricted to APCs, and has been previously linked to anti-mycobacterial immune responses in part through the induction of Th1/Th17-polarizing cytokines (43–46). While antigen-specific Th1 responses have long been considered to be necessary for natural and BCG-induced anti-TB immunity (64–66), recent work has described roles for non-canonical T cell subsets, including Th17 cells, in *Mtb* control (32,67) and in vaccine-induced protective responses (68–70) across both humans and animal models. hPC5 being enriched for Th17 differentiation and Th17-inducible cytokine signaling thus provides support for the idea that non-canonical T cell responses should be further considered as a species-translatable feature of BCG-mediated protective immunity. Moreover, evaluating the enrichment of these gene sets over time with ssGSEA (**Figs. 4D and S7**) reinforced our assertions that the human-relevant correlates of protection identified by our species translation model were BCG-induced and translatable even across different animal model species, albeit not time-agnostic.

Finally, we partially validated the protection-associated mechanisms inferred from our translation model using a publicly available microarray dataset collected from BCG-vaccinated South African infants which were categorized as cases or controls during two years of follow-up (35). The original study, which was unable to robustly characterize a common correlate of risk from the microarray data, instead identified two clusters of infants across follow-up outcomes by their patterns in gene expression. Of these two clusters, we found that infants belonging to Cluster 1 could be distinguished as cases or controls by their blood transcriptional expression of genes in the top 5% of (+) hPC12 by loadings. Expression of genes along either extreme of hPC5, which we also predicted to be associated with protection, did not significantly separate infants in this cohort with respect to their follow-up outcome. Partial validation of our hypotheses was not a surprising result, given that existing clinical data which attempts to characterize BCG-mediated protection outcomes are quite limited. Vaccination at birth, being standard-of-care in countries with high TB incidence, restricts the collection of data from non-vaccinated participants, which clouds the ability to distinguish between vaccine-induced and natural anti-TB immunity. Further, without deliberate pathogen exposure, it is possible that healthy controls comprise both protected and unexposed individuals. Moreover, as aptly demonstrated in Fletcher *et al* (35), variance among study participants can yield divergent host responses to vaccination and mask disease-relevant biological signals. Regardless of limitations in the validation data, we also acknowledge the caveat that our species translation modeling approach identifies human-relevant, i.e., conserved, signatures of protection from NHP data, and human relevance is not alone a sufficient condition for disease relevance in humans.

Overall, this study highlights the utility of translational cross-species modeling to leverage animal studies to uncover mechanisms of action in humans where no outcome data are available. While applied here to analyze human and NHP bulk blood transcriptomics to investigate BCG-mediated protective immunity, TransCompR is generalizable to a wide variety of biological contexts, species, and data modalities. Our computational framework is also well-suited for heterogeneous and highly variable datasets, facilitating a systems biology approach to identifying disease-relevant biological variance. Altogether, cross-species modeling is a useful tool to better understand the mechanisms underlying vaccine-mediated protective immunity, ultimately supporting the development of improved preventive interventions for TB.

## METHODS

### Cohorts and corresponding study designs

#### South African infant cohort

We performed a nested case-control study of participants from a parent cohort of 11,680 South African infants (6,7), who were vaccinated with BCG either percutaneously or intradermally within 24 hours of birth and from whom blood was collected at 10 weeks post-birth for immunological analyses (6). The vast majority (88.33%) of these infants were of mixed ethnic ancestry, with a median birth weight of 2.81 kg. In this work, we analyzed a group of 60 infants (29 male, 31 female) who had available cryopreserved PBMC and who had the highest or lowest IFNγ-expressing T cell responses to BCG, selected from 189 infants with data on BCG-specific cytokine responses (71). Selection of these infants was not informed by biological sex.

#### NHP dose cohort (GSE218270)

34 adult Indian-origin rhesus macaques (*M. mulatta*) (16 male, 18 female; median age 4.4 years) were included in the previously published dose cohort (33). Macaques were randomized into 6 vaccine groups to receive varying doses of intravenous BCG vaccination (4.5–7.5 log_10_ CFU, in half-log increments), and whole blood was collected four weeks prior to vaccination and at 2 days, 2 weeks, 4 weeks, and 12 weeks post-vaccination for bulk RNA-seq. Macaques were challenged 5–6 months after vaccination with a low dose of *Mtb* and euthanized 12 weeks later, or at the humane endpoint, for analysis of disease burden. As in the original publication, protection was defined as fewer than 100 total CFU *Mtb* in lung tissues upon necropsy. Processed sequencing data were obtained from GEO under accession number GSE218270. Corresponding flow cytometry and antibody titers measured at the same timepoints were obtained from [https://github.com/Khatri-Lab/bcg_transcriptome].

#### NHP route cohort (GSE218157)

35 adult Indian-origin rhesus macaques (*M. mulatta*) (17 male, 18 female; median age 4.7 years) were included in the previously published route cohort (36). Macaques received BCG vaccination through different routes which were randomly assigned: aerosol (n = 7), intradermal (n = 7), high-dose intradermal (n = 8), combined aerosol and intradermal (n = 7), and intravenous at 5 × 10^7^ CFU of BCG (n = 7). Whole blood was collected at baseline and at 2 days, 2 weeks, and 12 weeks post-vaccination for bulk RNA-seq. Macaques were later challenged with *Mtb* 6–10 months following BCG vaccination and euthanized 11–15 weeks following challenge or at humane endpoint for analysis of disease burden. As in the original publication, protection was defined as fewer than 100 total CFU *Mtb* [i.e., log_10_(CFU) < 2] in lung tissues upon necropsy. Processed sequencing data were obtained from GEO under accession number GSE218157.

#### Fletcher et al cohort (GSE20716)

46 BCG-vaccinated South African infants were included in the previously published microarray dataset from Fletcher *et al* (35). Blood was collected 10 weeks post-birth for PBMC isolation and microarray data analysis. Infants were then followed over a two-year period for characterization into confirmed TB cases (n = 26) or controls (n = 20). Processed microarray data were obtained from GEO under accession number GSE20716.

### Human blood collection, stimulation, and cryopreservation

At 10 weeks of age, heparinized venous blood was collected from all infants. For ICS, 1 ml whole blood was immediately incubated with BCG (SSI strain, 1.2 × 106 organisms/ml), as previously described (6). Whole blood incubated with medium alone served as negative control, and staphylococcal enterotoxin B (10 μg/ml; Sigma-Aldrich, St. Louis, MO) was used as positive control. The costimulatory antibodies, anti-CD28 and anti-CD49d (1 μg/ml each; BD Biosciences, San Jose, CA), were added to all conditions, for enhancement of specific responses. Blood was incubated for 7 hours at 37°C. Brefeldin-A was then added, followed by incubation for an additional 5 hours.

Cells were then harvested, fixed, and cryopreserved as previously described (72). Peripheral blood mononuclear cells (PBMCs) were isolated by Ficoll-density gradient centrifugation and cryopreserved in liquid nitrogen. For bulk RNA-seq, PBMCs were thawed, washed, and immediately used for RNA extraction.

### Human bulk RNA sequencing and analysis

RNA was isolated according to the manufacturer’s protocol using an miRNeasy Mini Kit (Qiagen). RNA integrity and yield was tested on the Agilent 2100 Bioanalyzer. The resulting cDNA was used to prepare a sequencing library. Libraries were sequenced on a MGI DNBSEQ-G400RS platform using DNBSEQ-G400RS high through-put sequencing set (FCL PE100), to acquire paired-end reads of 100 base pairs in length. Reads were aligned against the human reference genome: Homo sapiens GRCh38. Analysis of reads was performed in R v4.1.2. The data were log_2_-normalized (voom transformation) for downstream analyses. Lowly expressed genes were filtered by CPM greater than 2 in at least 5 samples.

### Human intracellular cytokine staining (ICS)

ICS was performed as previously described (71). Briefly, cryopreserved cells were thawed, washed, and permeabilized with Perm/wash solution (BD Biosciences). Cells were then incubated at 4°C for 1 hour with fluorescence-conjugated antibodies directed against surface antigens and intracellular cytokines. The following antibodies were used: anti-CD3 APC-H7 (clone SK7); anti-CD66 BV711 (B6.2/CD66); anti-CD14 PercP eFluor 710 (61D3); anti-CD33 BV650 (WM5); anti-CD16 AF488 (3G); anti-CD123 BV785 (6H6); anti-CD11c BV421 (3.9); anti–TNF-α Cy7PE (Mab11); anti–IFN-γ AF700 (B27); anti-IL-6 APC (MQ2-13A5); anti-PD-L1/CD274 PE (MIH1); anti-CD40 BV510 (5C3). Cells were acquired on a LSR II flow cytometer (BD Biosciences) configured with 3 lasers and 10 detectors, with FACS Diva 6.1 software (San Jose, CA). Optimal photomultiplier tube settings were established for this study before sample analysis.

Cytometer setting and tracking beads (BD Biosciences) were used to record the target median fluorescence intensity (MFI) values for the baseline settings, and these calibrations were performed each day before sample acquisition. Compensation settings were set with anti-mouse kappa-beads (BD Biosciences) labeled with the respective fluorochrome-conjugated antibodies. Flowjo version 8.8.4 (Treestar, Ashland, OR) was used to compensate and to analyze the flow cytometric data.

#### Regression of principal component scores onto covariates

Univariate regressions were performed to identify covariates that explain variation in score distributions on specific human-derived principal components. For binary covariates (human: responder group; NHP: binned post-challenge protection outcome), logistic regressions were performed, and z-values were reported as regression coefficients. For continuous covariates (human: ICS features; NHP: BCG dose, post-challenge lung bacterial burden), linear regressions were performed, and t-values were reported as regression coefficients. All regression coefficients are presented as averages ± SD from 5-fold cross-validation.

#### Translatable components regression (TransCompR)

Principal component analysis (PCA) was performed on the z-score-normalized, orthologue-limited human data using the prcomp function from the stats package (v3.6.2) in R to obtain the human sample scores and gene loadings on each human-derived principal component (hPC). The first 15 hPCs explained 80% of variance in the human data and were thus retained for downstream analyses. NHP samples were projected into this 15 hPC space by multiplying the z-score-normalized, orthologue-limited NHP data matrix by the human gene loadings matrix, resulting in a matrix of NHP scores in the hPC space. The variance of the NHP scores along all hPCs was totaled and used to normalize the amount of variance in the projected NHP data explained by each individual hPC. hPCs were then down-selected by whether their univariate regression coefficient was statistically significant across all 5 cross-validation folds. Univariate regression modeling was conducted using the glm function from the stats package (v3.6.2) in R.

#### TransCompR model validation

To ensure that we constructed a TransCompR model which predicted post-*Mtb* challenge bacterial burden with better performance than random chance, we compared prediction error or accuracy for covariate TransCompR models using all significantly univariately-associated hPCs to a series of null models with either random hPC selection or shuffled outcome variable (CFU or binned protection outcome). For all models, distributions of test error or accuracy scores were generated using 1000 trials of 5-fold cross-validation. Distributions were then statistically compared by Wilcoxon rank-sum testing.

#### Gene set enrichment analysis (GSEA)

To biologically interpret protection-associated hPCs, we used the Reactome Pathway (39) and KEGG (40) databases as references. For Reactome Pathway terms, we used the fgsea package (v1.21.1) in R (73) to calculate enrichment scores and estimate p-values. KEGG terms were limited to exclude those that were related to cancer or were specific to non-blood tissues. To calculate odds ratios for KEGG terms, we used the enrichR package (v3.3) in R (74), using the top 20% of genes along the extreme of each hPC as input. Single-sample GSEA (ssGSEA) was performed using select Reactome Pathway leading edge gene sets, and statistical testing between relevant groups (protected vs. non-protected) was conducted by unpaired t-test with Holm-Šídák’s multiple comparison correction across timepoints calculated in GraphPad.

#### Statistics

Summary data are presented as mean ± SD unless otherwise specified. Statistical analyses are specified in the figure legends. Values were generally considered significant at P < 0.05. All analyses were performed using either custom R scripts or GraphPad Prism (version 10.0.3).

#### Study approval

Human participants were enrolled from a parent study (6,7), which was conducted in accordance with the U.S. Department of Health and Human Services and Good Clinical Practice guidelines, and included protocol approval by the University of Cape Town Human Research Ethics Committee (332/2005) and written informed consent from the parent or legal guardian.

#### Data availability

The publicly available data utilized in this study can be found on NCBI GEO under the following accession numbers: GSE218270 (NHP dose cohort), GSE218157 (NHP route cohort), GSE20716 (infant microarray data). The BCG-vaccinated infant bulk RNA-seq published in this study are publicly available in Gene Expression Omnibus (GEO) at GSEXXXXXX (to be deposited upon publication). The TransCompR species translation modeling code was adapted from MathWorks File Exchange (number 77987).

## AUTHOR CONTRIBUTIONS

KB, CMS, TJS, and DAL conceived the study; DA, AG, and TDM performed experiments; KB, DA, AG, and TDM acquired and/or analyzed the data; KB wrote the original manuscript; CMS, TJS, and DAL supervised the study; and CMS, TJS, and DAL acquired funding. All authors reviewed, edited, and agreed to the final version of the manuscript.

## ACKNOWLEDGMENTS

This work was supported by NIH contract 75N93019C00071 to CMS and DAL; NIH grant AI181898 to CMS, TJS, and DAL; and the Army Institute for Collaborative Biotechnologies collaborative agreement W911NF-19-2-0026 to DAL. Work by KB was supported in part by the NIEHS Training Grant in Environmental Toxicology (Grant # T32-ES007020). The authors further thank Sarah Fortune, the HI-IMPAcTB consortium, and all members of the Lauffenburger lab for insightful discussions and advice.

## REFERENCES

1. Global Tuberculosis Report 2024. 1st ed. Geneva: World Health Organization; 2024. 1 p.

2. Davenne T, McShane H. Why don’t we have an effective tuberculosis vaccine yet? Expert Rev Vaccines. 2016 Aug 2;15(8):1009–13.

3. Martinez L, Cords O, Liu Q, Acuna-Villaorduna C, Bonnet M, Fox GJ, et al. Infant BCG vaccination and risk of pulmonary and extrapulmonary tuberculosis throughout the life course: a systematic review and individual participant data meta-analysis. The Lancet Global Health. 2022 Sep 1;10(9):e1307–16.

4. Roy A, Eisenhut M, Harris RJ, Rodrigues LC, Sridhar S, Habermann S, et al. Effect of BCG vaccination against Mycobacterium tuberculosis infection in children: systematic review and meta-analysis. BMJ. 2014 Aug 5;349:g4643.

5. Mangtani P, Abubakar I, Ariti C, Beynon R, Pimpin L, Fine PEM, et al. Protection by BCG Vaccine Against Tuberculosis: A Systematic Review of Randomized Controlled Trials. Clinical Infectious Diseases. 2014 Feb 15;58(4):470–80.

6. Kagina BMN, Abel B, Scriba TJ, Hughes EJ, Keyser A, Soares A, et al. Specific T Cell Frequency and Cytokine Expression Profile Do Not Correlate with Protection against Tuberculosis after Bacillus Calmette-Guérin Vaccination of Newborns. Am J Respir Crit Care Med. 2010 Oct 15;182(8):1073–9.

7. Hawkridge A, Hatherill M, Little F, Goetz MA, Barker L, Mahomed H, et al. Efficacy of percutaneous versus intradermal BCG in the prevention of tuberculosis in South African infants: randomised trial. BMJ. 2008 Nov 13;337:a2052.

8. Zak DE, Penn-Nicholson A, Scriba TJ, Thompson E, Suliman S, Amon LM, et al. A blood RNA signature for tuberculosis disease risk: a prospective cohort study. The Lancet. 2016 Jun 4;387(10035):2312–22.

9. Huang Y, Ai L, Wang X, Sun Z, Wang F. Review and Updates on the Diagnosis of Tuberculosis. Journal of Clinical Medicine. 2022 Jan;11(19):5826.

10. MacGregor-Fairlie M, Wilkinson S, Besra GS, Goldberg Oppenheimer P. Tuberculosis diagnostics: overcoming ancient challenges with modern solutions. Emerg Top Life Sci. 2020 Dec 11;4(4):435– 48.

11. Chopra KK, Singh S. Newer diagnostic tests for tuberculosis, their utility, and their limitations. Current Medicine Research and Practice. 2020 Jan 1;10(1):8–11.

12. Wallis RS, Doherty TM, Onyebujoh P, Vahedi M, Laang H, Olesen O, et al. Biomarkers for tuberculosis disease activity, cure, and relapse. Lancet Infect Dis. 2009 Mar;9(3):162–72.

13. Doherty M, Wallis RS, Zumla A, WHO-Tropical Disease Research/European Commission joint expert consultation group. Biomarkers for tuberculosis disease status and diagnosis. Curr Opin Pulm Med. 2009 May;15(3):181–7.

14. Russell DG, Barry CE, Flynn JL. Tuberculosis: What We Don’t Know Can, and Does, Hurt Us. Science. 2010 May 14;328(5980):852–6.

15. Li H, Li H. Animal Models of Tuberculosis. In: Christodoulides M, editor. Vaccines for Neglected Pathogens: Strategies, Achievements and Challenges: Focus on Leprosy, Leishmaniasis, Melioidosis and Tuberculosis [Internet]. Cham: Springer International Publishing; 2023 [cited 2025 Mar 18]. p. 139–70. Available from: 10.1007/978-3-031-24355-4_7

16. Singh AK, Gupta UD. Animal models of tuberculosis: Lesson learnt. Indian J Med Res. 2018 May;147(5):456–63.

17. Seyhan AA. Lost in translation: the valley of death across preclinical and clinical divide – identification of problems and overcoming obstacles. Translational Medicine Communications. 2019 Nov 18;4(1):18.

18. Hackam DG, Redelmeier DA. Translation of research evidence from animals to humans. JAMA. 2006 Oct 11;296(14):1731–2.

19. Shay T, Jojic V, Zuk O, Rothamel K, Puyraimond-Zemmour D, Feng T, et al. Conservation and divergence in the transcriptional programs of the human and mouse immune systems. Proceedings of the National Academy of Sciences. 2013 Feb 19;110(8):2946–51.

20. Godec J, Tan Y, Liberzon A, Tamayo P, Bhattacharya S, Butte AJ, et al. Compendium of Immune Signatures Identifies Conserved and Species-Specific Biology in Response to Inflammation. Immunity. 2016 Jan 19;44(1):194–206.

21. Pullen KM, Finethy R, Ko SHB, Reames CJ, Sassetti CM, Lauffenburger DA. Cross-species transcriptomics translation reveals a role for the unfolded protein response in Mycobacterium tuberculosis infection. npj Syst Biol Appl. 2025 Feb 15;11(1):1–13.

22. Abebe F. Is interferon-gamma the right marker for bacille Calmette–Guérin-induced immune protection? The missing link in our understanding of tuberculosis immunology. Clin Exp Immunol. 2012 Sep;169(3):213–9.

23. Andersen P, Smedegaard B. CD4(+) T-cell subsets that mediate immunological memory to Mycobacterium tuberculosis infection in mice. Infect Immun. 2000 Feb;68(2):621–9.

24. Caruso AM, Serbina N, Klein E, Triebold K, Bloom BR, Flynn JL. Mice deficient in CD4 T cells have only transiently diminished levels of IFN-gamma, yet succumb to tuberculosis. J Immunol. 1999 May 1;162(9):5407–16.

25. Soares AP, Scriba TJ, Joseph S, Harbacheuski R, Murray RA, Gelderbloem SJ, et al. Bacille Calmette Guerin vaccination of human newborns induces T cells with complex cytokine and phenotypic profiles. J Immunol. 2008 Mar 1;180(5):3569–77.

26. Normand R, Du W, Briller M, Gaujoux R, Starosvetsky E, Ziv-Kenet A, et al. Found In Translation: a machine learning model for mouse-to-human inference. Nat Methods. 2018 Dec;15(12):1067–73.

27. Brubaker DK, Kumar MP, Chiswick EL, Gregg C, Starchenko A, Vega PN, et al. An interspecies translation model implicates integrin signaling in infliximab-resistant inflammatory bowel disease. Sci Signal. 2020 Aug 4;13(643):eaay3258.

28. Kowald A, Barrantes I, Möller S, Palmer D, Murua Escobar H, Schwerk A, et al. Transfer learning of clinical outcomes from preclinical molecular data, principles and perspectives. Brief Bioinform. 2022 May 13;23(3):bbac133.

29. Suarez-Lopez L, Shui B, Brubaker DK, Hill M, Bergendorf A, Changelian PS, et al. Cross-species transcriptomic signatures predict response to MK2 inhibition in mouse models of chronic inflammation. iScience. 2021 Dec 17;24(12):103406.

30. Lee MJ, Wang C, Carroll MJ, Brubaker DK, Hyman BT, Lauffenburger DA. Computational Interspecies Translation Between Alzheimer’s Disease Mouse Models and Human Subjects Identifies Innate Immune Complement, TYROBP, and TAM Receptor Agonist Signatures, Distinct From Influences of Aging. Front Neurosci [Internet]. 2021 Sep 30 [cited 2025 Jan 21];15. Available from: https://www.frontiersin.org/journals/neuroscience/articles/10.3389/fnins.2021.727784/full

31. Carroll MJ, Garcia-Reyero N, Perkins EJ, Lauffenburger DA. Translatable pathways classification (TransPath-C) for inferring processes germane to human biology from animal studies data: example application in neurobiology. Integr Biol (Camb). 2021 Dec 15;13(10):237–45.

32. Proulx MK, Wiggins CD, Reames CJ, Wu C, Kiritsy MC, Xu P, et al. Noncanonical T cell responses are associated with protection from tuberculosis in mice and humans. Journal of Experimental Medicine. 2025 Apr 7;222(7):e20241760.

33. Liu YE, Darrah PA, Zeppa JJ, Kamath M, Laboune F, Douek DC, et al. Blood transcriptional correlates of BCG-induced protection against tuberculosis in rhesus macaques. Cell Rep Med. 2023 Jul 18;4(7):101096.

34. Darrah PA, Zeppa JJ, Maiello P, Hackney JA, Wadsworth MH, Hughes TK, et al. Prevention of tuberculosis in macaques after intravenous BCG immunization. Nature. 2020 Jan;577(7788):95–102.

35. Fletcher HA, Filali-Mouhim A, Nemes E, Hawkridge A, Keyser A, Njikan S, et al. Human newborn bacille Calmette–Guérin vaccination and risk of tuberculosis disease: a case-control study. BMC Medicine. 2016 May 16;14(1):76.

36. Darrah PA, Zeppa JJ, Wang C, Irvine EB, Bucsan AN, Rodgers MA, et al. Airway T cells are a correlate of i.v. Bacille Calmette-Guerin-mediated protection against tuberculosis in rhesus macaques. Cell Host Microbe. 2023 Jun 14;31(6):962–977.e8.

37. Irvine EB, Darrah PA, Wang S, Wang C, McNamara RP, Roederer M, et al. Humoral correlates of protection against Mycobacterium tuberculosis following intravenous BCG vaccination in rhesus macaques. iScience [Internet]. 2024 Dec 20 [cited 2025 Feb 5];27(12). Available from: https://www.cell.com/iscience/abstract/S2589-0042(24)02353-8

38. Meimetis N, Pullen KM, Zhu DY, Nilsson A, Hoang TN, Magliacane S, et al. AutoTransOP: translating omics signatures without orthologue requirements using deep learning. npj Syst Biol Appl. 2024 Jan 29;10(1):1–19.

39. Milacic M, Beavers D, Conley P, Gong C, Gillespie M, Griss J, et al. The Reactome Pathway Knowledgebase 2024. Nucleic Acids Research. 2024 Jan 5;52(D1):D672–8.

40. Kanehisa M, Furumichi M, Sato Y, Matsuura Y, Ishiguro-Watanabe M. KEGG: biological systems database as a model of the real world. Nucleic Acids Res. 2025 Jan 6;53(D1):D672–7.

41. Ahmed M, Thirunavukkarasu S, Rosa BA, Thomas KA, Das S, Rangel-Moreno J, et al. Immune correlates of tuberculosis disease and risk translate across species. Science Translational Medicine. 2020 Jan 29;12(528):eaay0233.

42. Rosa BA, Ahmed M, Singh DK, Choreño-Parra JA, Cole J, Jiménez-Álvarez LA, et al. IFN signaling and neutrophil degranulation transcriptional signatures are induced during SARS-CoV-2 infection. Commun Biol. 2021 Mar 5;4(1):1–14.

43. Mata-Martínez P, Bergón-Gutiérrez M, del Fresno C. Dectin-1 Signaling Update: New Perspectives for Trained Immunity. Front Immunol. 2022 Feb 14;13:812148.

44. Rothfuchs AG, Bafica A, Feng CG, Egen JG, Williams DL, Brown GD, et al. Dectin-1 Interaction with Mycobacterium tuberculosis Leads to Enhanced IL-12p40 Production by Splenic Dendritic Cells1. The Journal of Immunology. 2007 Sep 15;179(6):3463–71.

45. Yadav M, Schorey JS. The β-glucan receptor dectin-1 functions together with TLR2 to mediate macrophage activation by mycobacteria. Blood. 2006 Nov 1;108(9):3168–75.

46. Wagener M, Hoving JC, Ndlovu H, Marakalala MJ. Dectin-1-Syk-CARD9 Signaling Pathway in TB Immunity. Front Immunol [Internet]. 2018 Feb 13 [cited 2025 Jan 23];9. Available from: https://www.frontiersin.org/journals/immunology/articles/10.3389/fimmu.2018.00225/full

47. Zhu X, Zhu J. CD4 T Helper Cell Subsets and Related Human Immunological Disorders. Int J Mol Sci. 2020 Oct 28;21(21):8011.

48. Pai M, Nicol MP, Boehme CC. Tuberculosis Diagnostics: State of the Art and Future Directions. Microbiology Spectrum. 2016 Oct 21;4(5):10.1128/microbiolspec.tbtb2-0019–2016.

49. Fletcher HA, Snowden MA, Landry B, Rida W, Satti I, Harris SA, et al. T-cell activation is an immune correlate of risk in BCG vaccinated infants. Nat Commun. 2016 Apr 12;7(1):11290.

50. Mittrücker HW, Steinhoff U, Köhler A, Krause M, Lazar D, Mex P, et al. Poor correlation between BCG vaccination-induced T cell responses and protection against tuberculosis. Proceedings of the National Academy of Sciences. 2007 Jul 24;104(30):12434–9.

51. Smith SG, Zelmer A, Blitz R, Fletcher HA, Dockrell HM. Polyfunctional CD4 T-cells correlate with *in vitro* mycobacterial growth inhibition following *Mycobacterium bovis* BCG-vaccination of infants. Vaccine. 2016 Oct 17;34(44):5298–305.

52. Bizzell E, Sia JK, Quezada M, Enriquez A, Georgieva M, Rengarajan J. Deletion of BCG Hip1 protease enhances dendritic cell and CD4 T cell responses. J Leukoc Biol. 2018 Apr;103(4):739–48.

53. Su H, Peng B, Zhang Z, Liu Z, Zhang Z. The *Mycobacterium tuberculosis* glycoprotein Rv1016c protein inhibits dendritic cell maturation, and impairs Th1 /Th17 responses during mycobacteria infection. Molecular Immunology. 2019 May 1;109:58–70.

54. Kleinnijenhuis J, Quintin J, Preijers F, Joosten LAB, Ifrim DC, Saeed S, et al. Bacille Calmette-Guerin induces NOD2-dependent nonspecific protection from reinfection via epigenetic reprogramming of monocytes. Proc Natl Acad Sci U S A. 2012 Oct 23;109(43):17537–42.

55. Moorlag SJCFM, Folkman L, ter Horst R, Krausgruber T, Barreca D, Schuster LC, et al. Multi-omics analysis of innate and adaptive responses to BCG vaccination reveals epigenetic cell states that predict trained immunity. Immunity. 2024 Jan 9;57(1):171–187.e14.

56. Li W, Moorlag SJCFM, Koeken VACM, Röring RJ, de Bree LCJ, Mourits VP, et al. A single-cell view on host immune transcriptional response to in vivo BCG-induced trained immunity. Cell Rep. 2023 May 30;42(5):112487.

57. Sun SJ, Aguirre-Gamboa R, de Bree LCJ, Sanz J, Dumaine A, van der Velden WJFM, et al. BCG vaccination alters the epigenetic landscape of progenitor cells in human bone marrow to influence innate immune responses. Immunity. 2024 Sep 10;57(9):2095–2107.e8.

58. Jeyanathan M, Vaseghi-Shanjani M, Afkhami S, Grondin JA, Kang A, D’Agostino MR, et al. Parenteral BCG vaccine induces lung-resident memory macrophages and trained immunity via the gut–lung axis. Nat Immunol. 2022 Dec;23(12):1687–702.

59. Hole CR, Wager CML, Castro-Lopez N, Campuzano A, Cai H, Wozniak KL, et al. Induction of memory-like dendritic cell responses in vivo. Nat Commun. 2019 Jul 4;10(1):2955.

60. Khan A, Singh VK, Mishra A, Soudani E, Bakhru P, Singh CR, et al. NOD2/RIG-I Activating Inarigivir Adjuvant Enhances the Efficacy of BCG Vaccine Against Tuberculosis in Mice. Front Immunol [Internet]. 2020 Dec 7 [cited 2025 Apr 7];11. Available from: https://www.frontiersin.org/journals/immunology/articles/10.3389/fimmu.2020.592333/full

61. Satti I, Wittenberg RE, Li S, Harris SA, Tanner R, Cizmeci D, et al. Inflammation and immune activation are associated with risk of Mycobacterium tuberculosis infection in BCG-vaccinated infants. Nat Commun. 2022 Nov 3;13(1):6594.

62. Donovan ML, Schultz TE, Duke TJ, Blumenthal A. Type I Interferons in the Pathogenesis of Tuberculosis: Molecular Drivers and Immunological Consequences. Front Immunol. 2017 Nov 27;8:1633.

63. Moreira-Teixeira L, Mayer-Barber K, Sher A, O’Garra A. Type I interferons in tuberculosis: Foe and occasionally friend. J Exp Med. 2018 May 7;215(5):1273–85.

64. Bustamante J, Boisson-Dupuis S, Abel L, Casanova JL. Mendelian susceptibility to mycobacterial disease: genetic, immunological, and clinical features of inborn errors of IFN-γ immunity. Semin Immunol. 2014 Dec;26(6):454–70.

65. Boisson-Dupuis S, Ramirez-Alejo N, Li Z, Patin E, Rao G, Kerner G, et al. Tuberculosis and impaired IL-23–dependent IFN-γ immunity in humans homozygous for a common TYK2 missense variant. Science Immunology. 2018 Dec 21;3(30):eaau8714.

66. Marchant A, Goetghebuer T, Ota MO, Wolfe I, Ceesay SJ, De Groote D, et al. Newborns develop a Th1-type immune response to Mycobacterium bovis bacillus Calmette-Guérin vaccination. J Immunol. 1999 Aug 15;163(4):2249–55.

67. Sun M, Phan JM, Kieswetter NS, Huang H, Yu KKQ, Smith MT, et al. Specific CD4+ T cell phenotypes associate with bacterial control in people who “resist” infection with Mycobacterium tuberculosis. Nat Immunol. 2024 Aug;25(8):1411–21.

68. Singhania A, Dubelko P, Kuan R, Chronister WD, Muskat K, Das J, et al. CD4+CCR6+ T cells dominate the BCG-induced transcriptional signature. eBioMedicine [Internet]. 2021 Dec 1 [cited 2025 Apr 8];74. Available from: https://www.thelancet.com/journals/ebiom/article/PIIS2352-3964(21)00540-5/fulltext

69. Wozniak TM, Saunders BM, Ryan AA, Britton WJ. Mycobacterium bovis BCG-Specific Th17 Cells Confer Partial Protection against Mycobacterium tuberculosis Infection in the Absence of Gamma Interferon. Infection and Immunity. 2010 Oct;78(10):4187–94.

70. Gopal R, Lin Y, Obermajer N, Slight S, Nuthalapati N, Ahmed M, et al. IL-23-dependent IL-17 drives Th1-cell responses following Mycobacterium bovis BCG vaccination. Eur J Immunol. 2012 Feb;42(2):364–73.

71. Anterasian C, Gela A, Mwambene TD, Shah JA, Ivie J, Dill-McFarland KA, et al. BCG induced innate immune response heterogeneity and susceptibility to pediatric tuberculosis. The Journal of Immunology. 2025 Apr 2;vkae062.

72. Hanekom WA, Hughes J, Mavinkurve M, Mendillo M, Watkins M, Gamieldien H, et al. Novel application of a whole blood intracellular cytokine detection assay to quantitate specific T-cell frequency in field studies. J Immunol Methods. 2004 Aug;291(1–2):185–95.

73. Korotkevich G, Sukhov V, Budin N, Shpak B, Artyomov MN, Sergushichev A. Fast gene set enrichment analysis [Internet]. bioRxiv; 2021 [cited 2025 Apr 10]. p. 060012. Available from: https://www.biorxiv.org/content/10.1101/060012v3

74. Kuleshov MV, Jones MR, Rouillard AD, Fernandez NF, Duan Q, Wang Z, et al. Enrichr: a comprehensive gene set enrichment analysis web server 2016 update. Nucleic Acids Res. 2016 Jul 8;44(W1):W90–97.

