## Supplemental Figs 1-7 for "Cross-species blood transcriptional correlates of BCG-mediated protection against tuberculosis include innate and adaptive immune processes"

*Figure S1: ICS provides additional context for human-derived PCs*

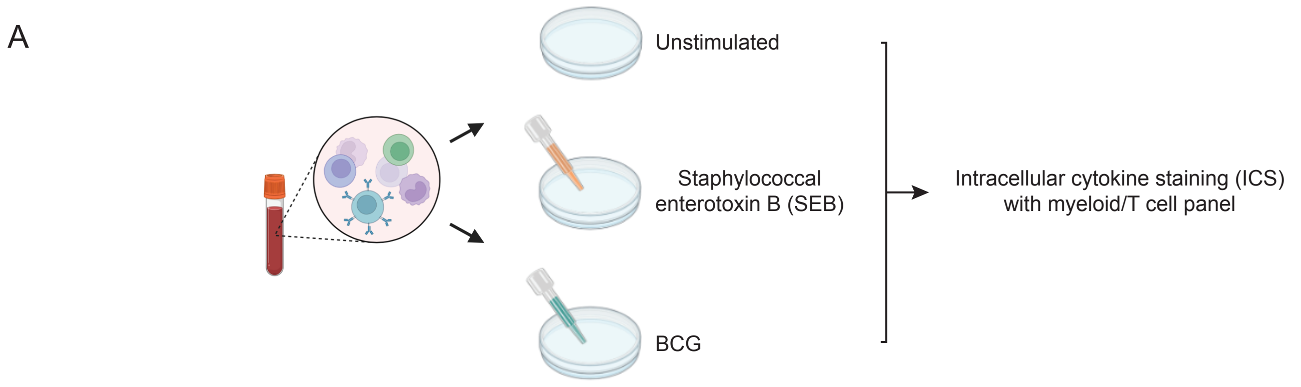


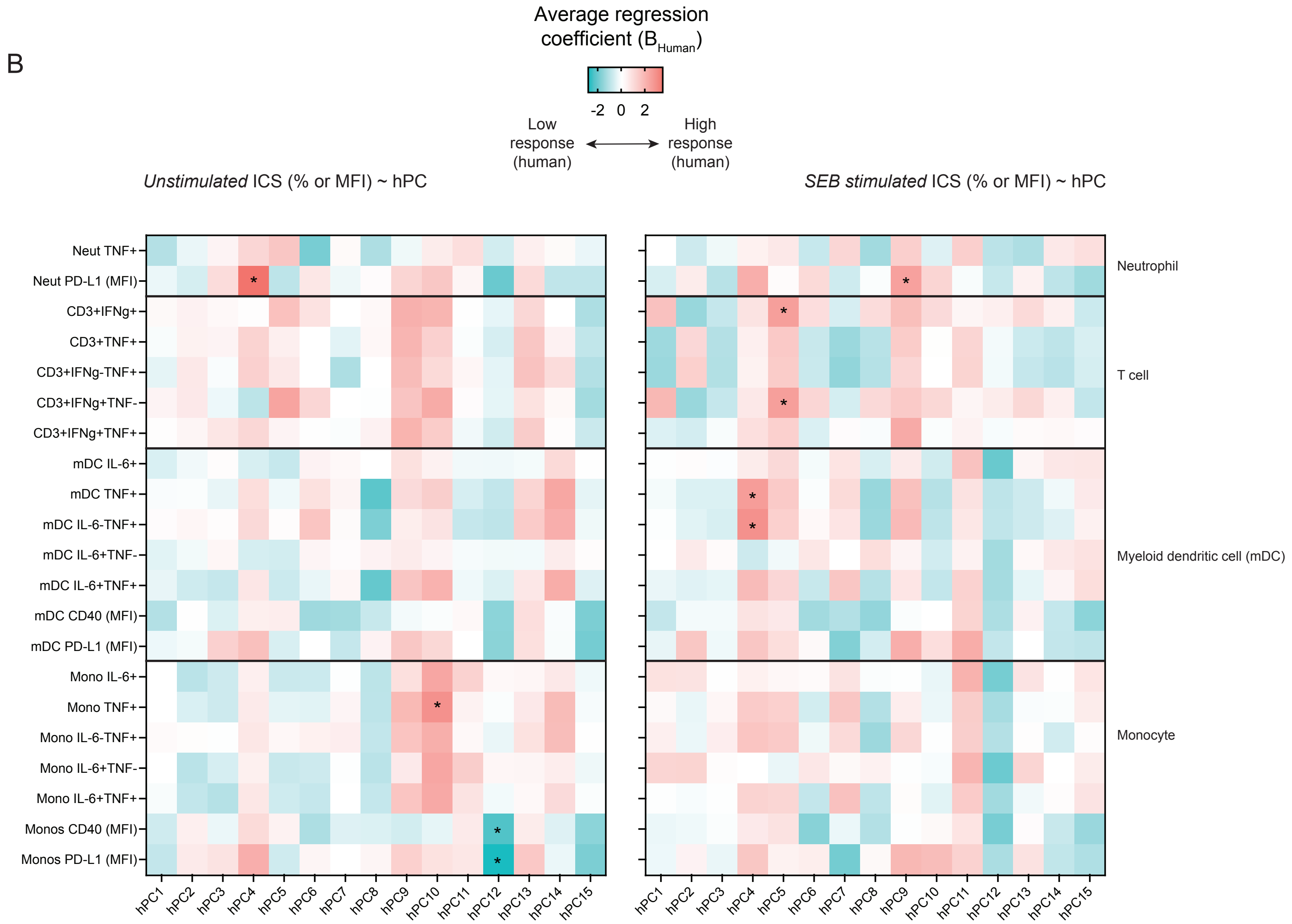


Related to Figs. 1 and 2.
(A) Schematic detailing collection of intracellular cytokine staining (ICS) data from infants profiled for bulk RNA-seq. SEB stimulation was used as a positive control to test for non-BCG-specific immune responses.
(B) Heatmaps detailing univariate regression coefficients from individual human-derived principal components regressed against ICS panel readouts (% unless otherwise indicated; y-axis) from unstimulated (left) and SEB-stimulated (right) *ex vivo* conditions. Regression coefficients are averaged across results from 5-fold cross validation. *p < 0.05 across folds.

*Figure S2: hPCs can separate human samples by BCG-specific Th1 response*

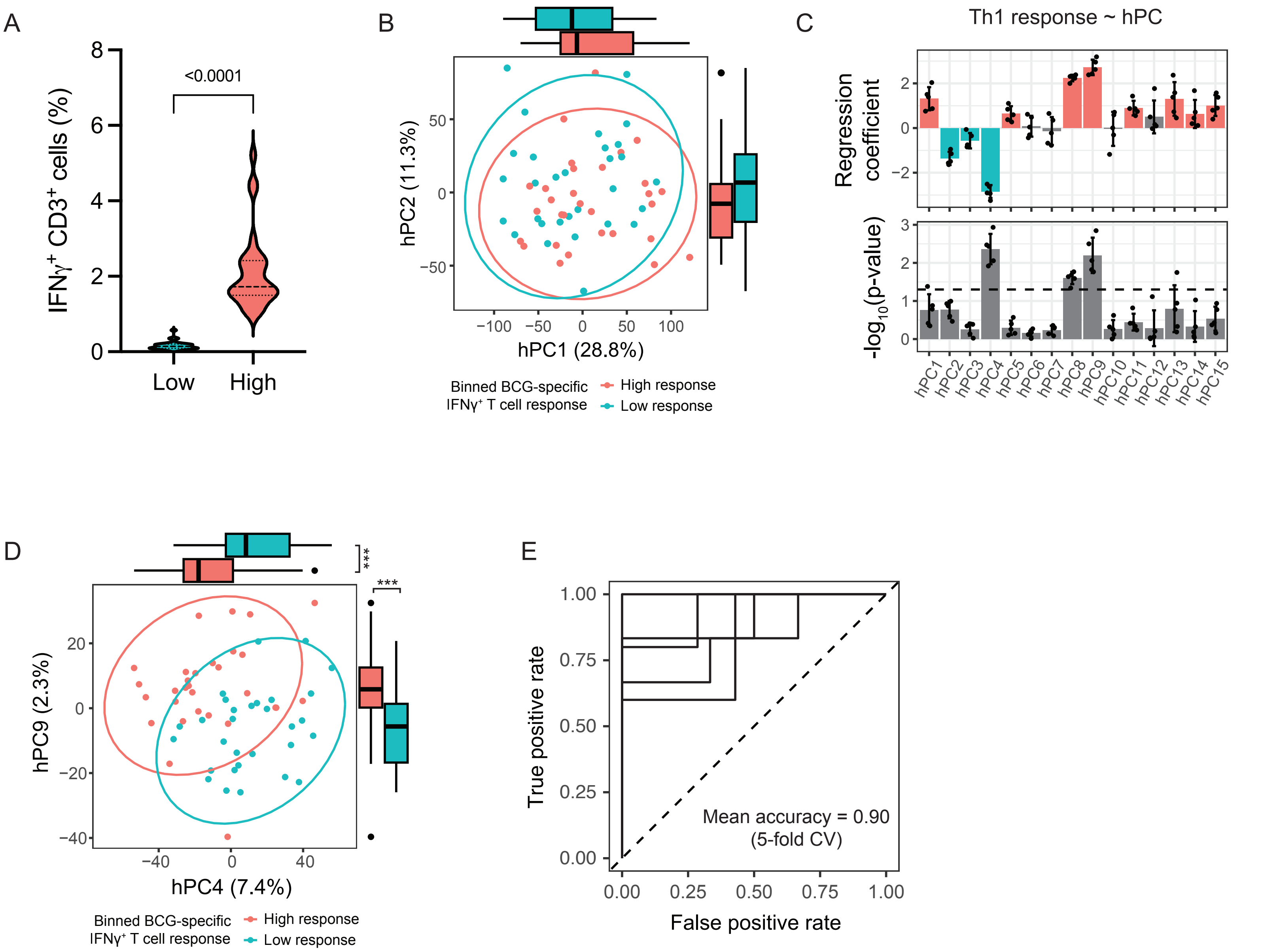


Related to Fig. 2.
(A) Percentage of CD3⁺ cells (T cells) expressing IFNγ upon *ex vivo* BCG stimulation separated into low (blue) and high (pink) responders. The dotted lines illustrate median and 1st and 3rd quartiles. P-value < 0.0001 by unpaired t-test as calculated in GraphPad.
(B and D) Human samples projected along (B) hPC1 and hPC2, and (D) hPC4 and hPC9. Each datapoint represents one sample, and the colored ellipses represent the 95% confidence interval for each phenotype group’s distribution in each respective space. Boxplots on the x- and y-axes illustrate the distribution of the high (pink) and low (blue) BCG-specific T cell response samples on each axis respectively. Thick lines denote the median; hinges denotes 1st and 3rd quartiles; whiskers extend from hinges ± 1.5 * IQR; outliers are plotted individually. ***p<0.001 by Wilcoxon rank-sum testing.
(C) (top) Bar plot showing 5-fold logistic regression coefficients (summary data presented as mean ± SD) for each hPC when univariately regressed against human phenotype (BCG response). (bottom) Bar plot showing the distribution of corresponding p-values.
(E) ROC curves for 5-fold logisitic regression of human phenotypes using projected human data onto hPC4, hPC8, and hPC9 together.

*Figure S3: Interpretation of hPC-projected NHP data w.r.t. BCG vaccine dose*

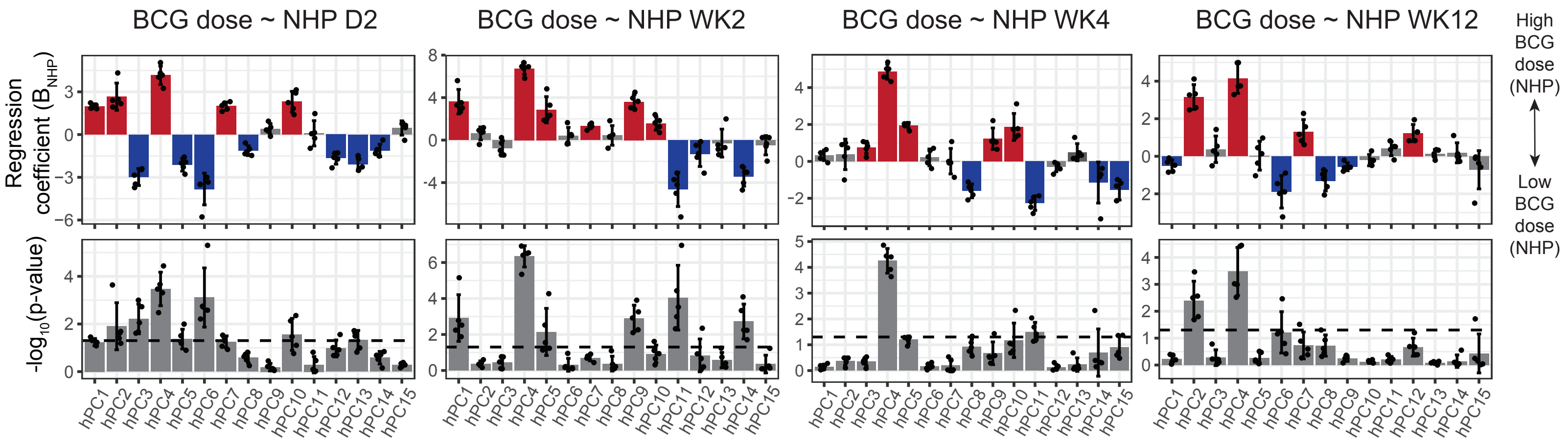


Related to Fig. 3.

Bar graphs showing (top) 5-fold logistic regression coefficients (summary data presented as mean ± SD) and (bottom) distribution of corresponding p-values from univariate regression of each hPC against BCG dose. The dotted line denotes p = 0.05.

*Figure S4: hPCs can reliably separate NHP samples by protection status, although model performance is dependent on time post-vaccination*

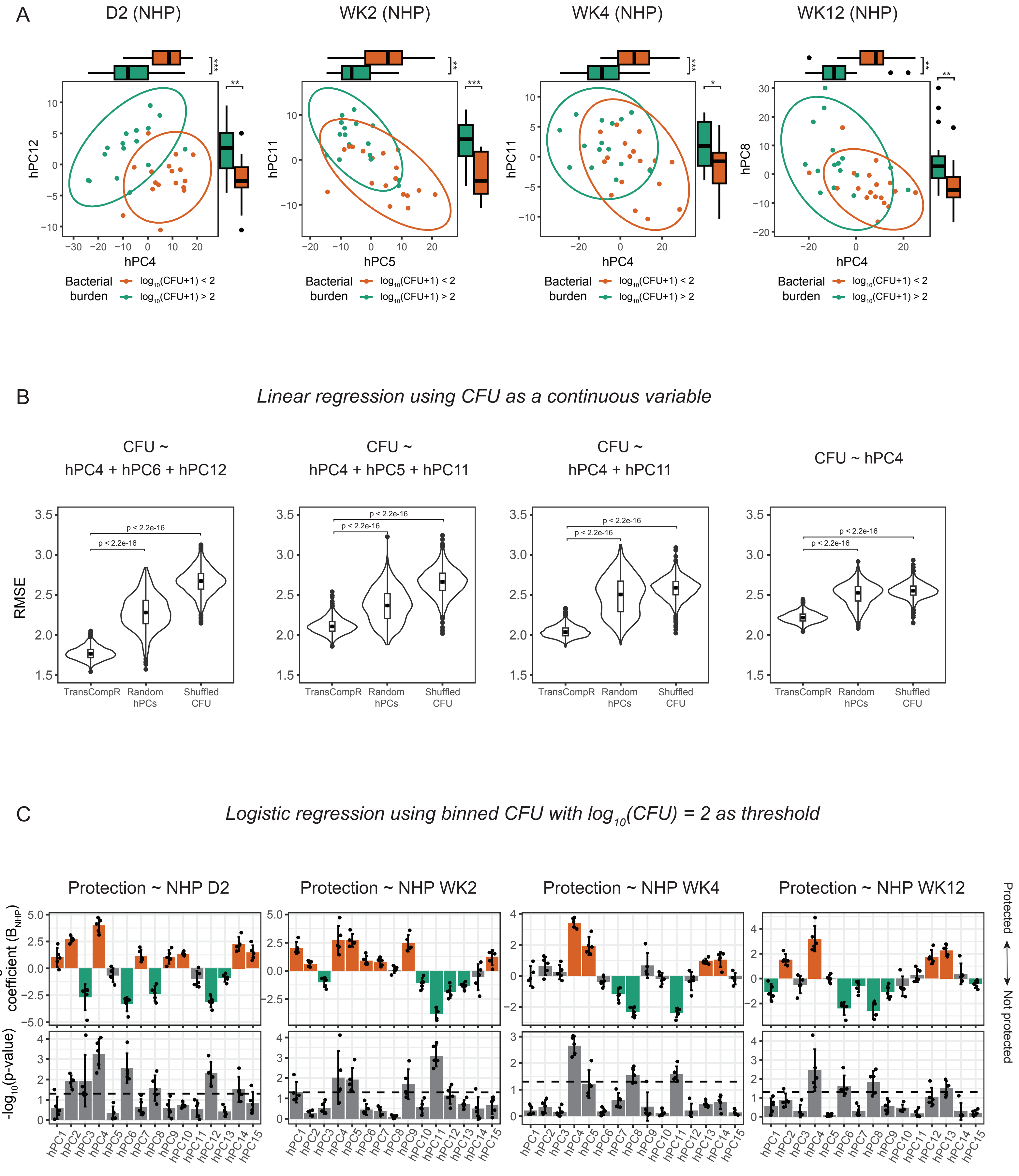

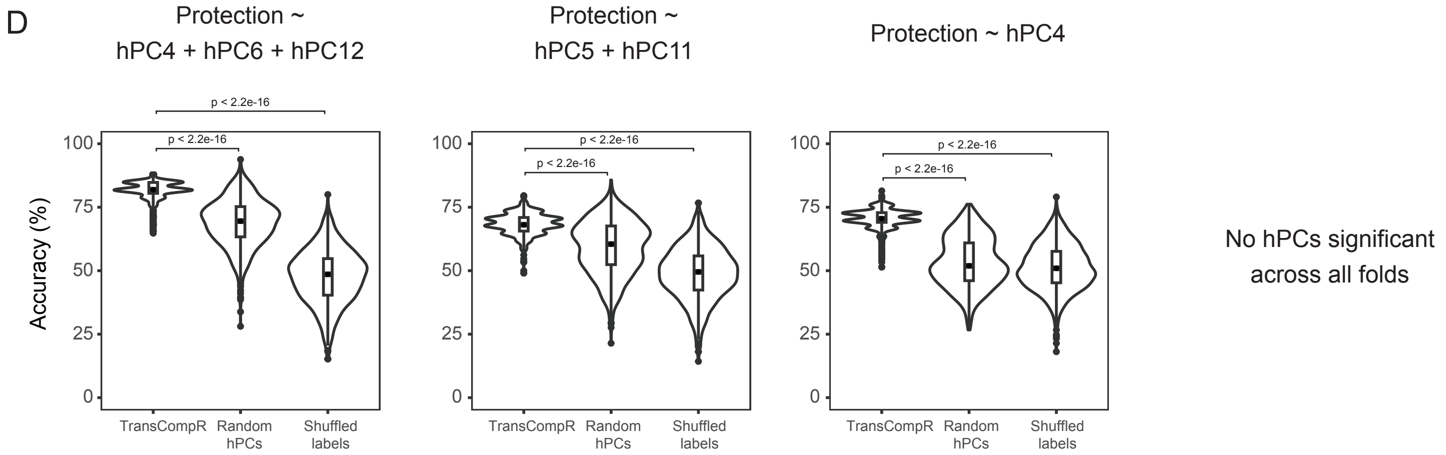


Related to Fig. 3.
(A) NHP samples from 4 post-vaccination timepoints projected along hPC4 and hPC12, hPC5 and hPC11, hPC4 and hPC11, and hPC4 and hPC8, respectively. Each datapoint represents one sample, and the colored ellipses represent the 95% confidence interval for each phenotype group's distribution in each respective space. Boxplots on the x- and y-axes illustrate the distribution of the high (green) and low (orange) CFU samples on each axis respectively. Thick lines denote the median; hinges denotes 1st and 3rd quartiles; whiskers extend from hinges ± 1.5 * IQR; outliers are plotted individually. *p<0.05, **p<0.01, ***p<0.001 by Wilcoxon rank-sum testing.
(B) Violin plots summarizing the test root mean square error (RMSE) for 1000 trials of 5-fold CV for covariate linear regression models constructed using (left) all hPCs that were statistically significant via univariate linear regression against numerical CFU (center) randomly selected hPCs, or (right) statistically significant hPCs and regressed against shuffled CFU. Boxplots within the violins are rendered as in (A). p < 2.2e-16 by Wilcoxon rank-sum testing.
(C) Bar graphs showing (top) 5-fold logistic regression coefficients (summary data presented as mean ± SD) and (bottom) distribution of corresponding p-values from univariate regression of each hPC against binned CFU (protected vs. not protected). The dotted line denotes p = 0.05.
(D) Violin plots summarizing the test accuracy (%) for 1000 trials of 5-fold CV for covariate logistic regression models constructed using (left) all hPCs that were statistically significant via univariate logistic regression against binned CFU in (C), (center) randomly selected hPCs, or (right) statistically significant hPCs and regressed against shuffled labels. Boxplots within the violins are rendered as in (A-B).p < 2.2e-16 by Wilcoxon rank-sum testing.

*Figure S5: Human-to-NHP translation with route cohort further implicates two protection-associated hPCs*

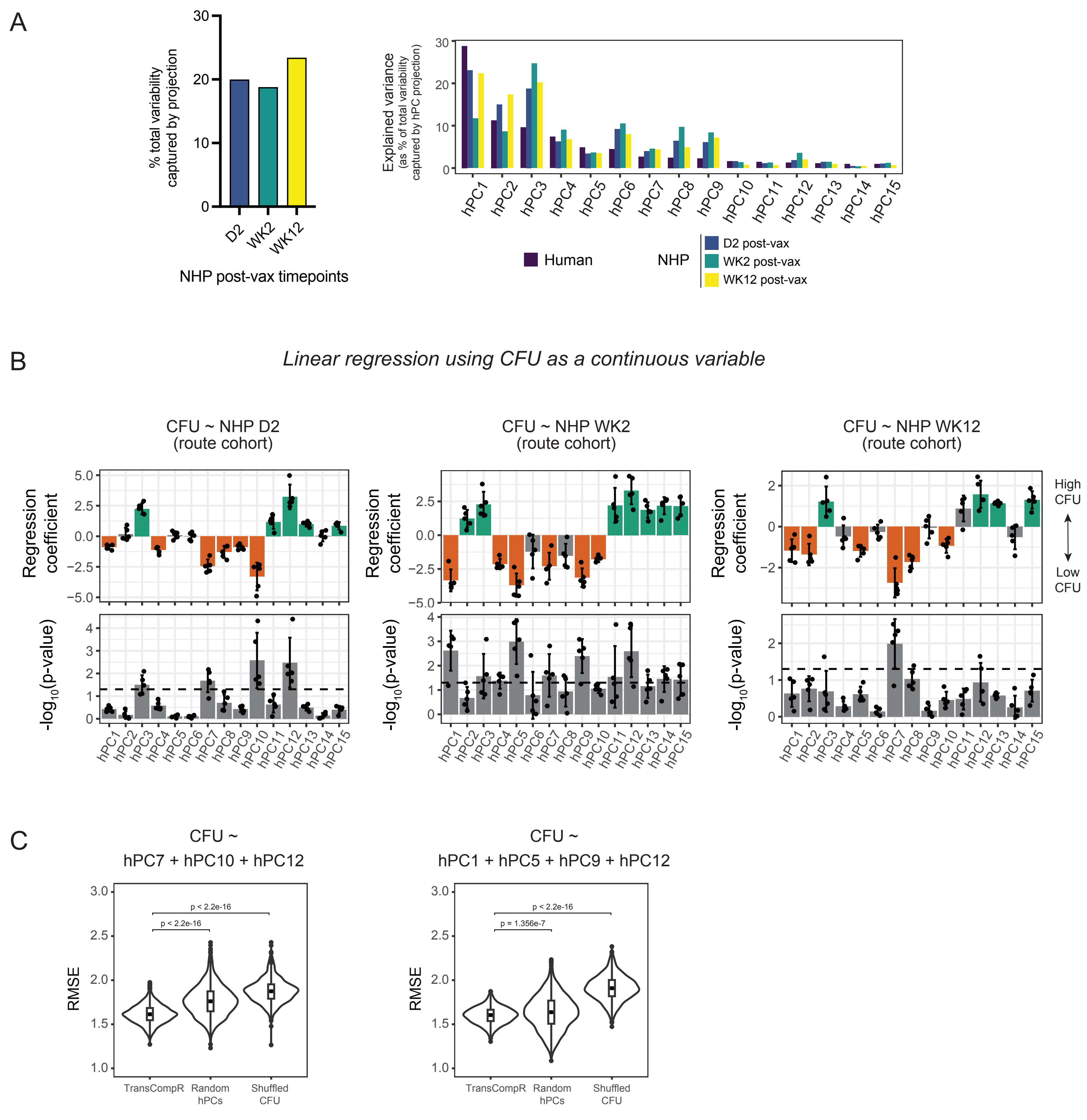


Related to Fig. 3.

(A) Bar graphs showing explained variance captured by projecting the NHP route cohort (GSE218157) into hPC space (left) as a percentage of total variance in each post-vaccination NHP dataset and (right) as distributed across each hPC in the TransCompR model.

(B) Bar graphs showing (top) 5-fold logistic regression coefficients (summary data presented as mean ± SD) and (bottom) distribution of corresponding p-values from univariate regression of each hPC against post-*Mtb* challenge bacterial burden (CFU). The dotted line denotes p = 0.05.

(C) Violin plots summarizing the test root mean square error (RMSE) for 1000 trials of 5-fold CV for covariate linear regression models constructed using (left) all hPCs that were statistically significant via univariate linear regression against numerical CFU in (B), (center) randomly selected hPCs, or (right) statistically significant hPCs and regressed against shuffled CFU. For boxplots within the violins: thick lines denote the median; hinges denotes 1st and 3rd quartiles; whiskers extend from hinges ± 1.5 * IQR; outliers are plotted individually. $p < 2.2e-16$ by Wilcoxon rank-sum testing.

*Figure S6: GSEA using KEGG database as reference provides additional support for biological interpretation of protection-associated hPCs*

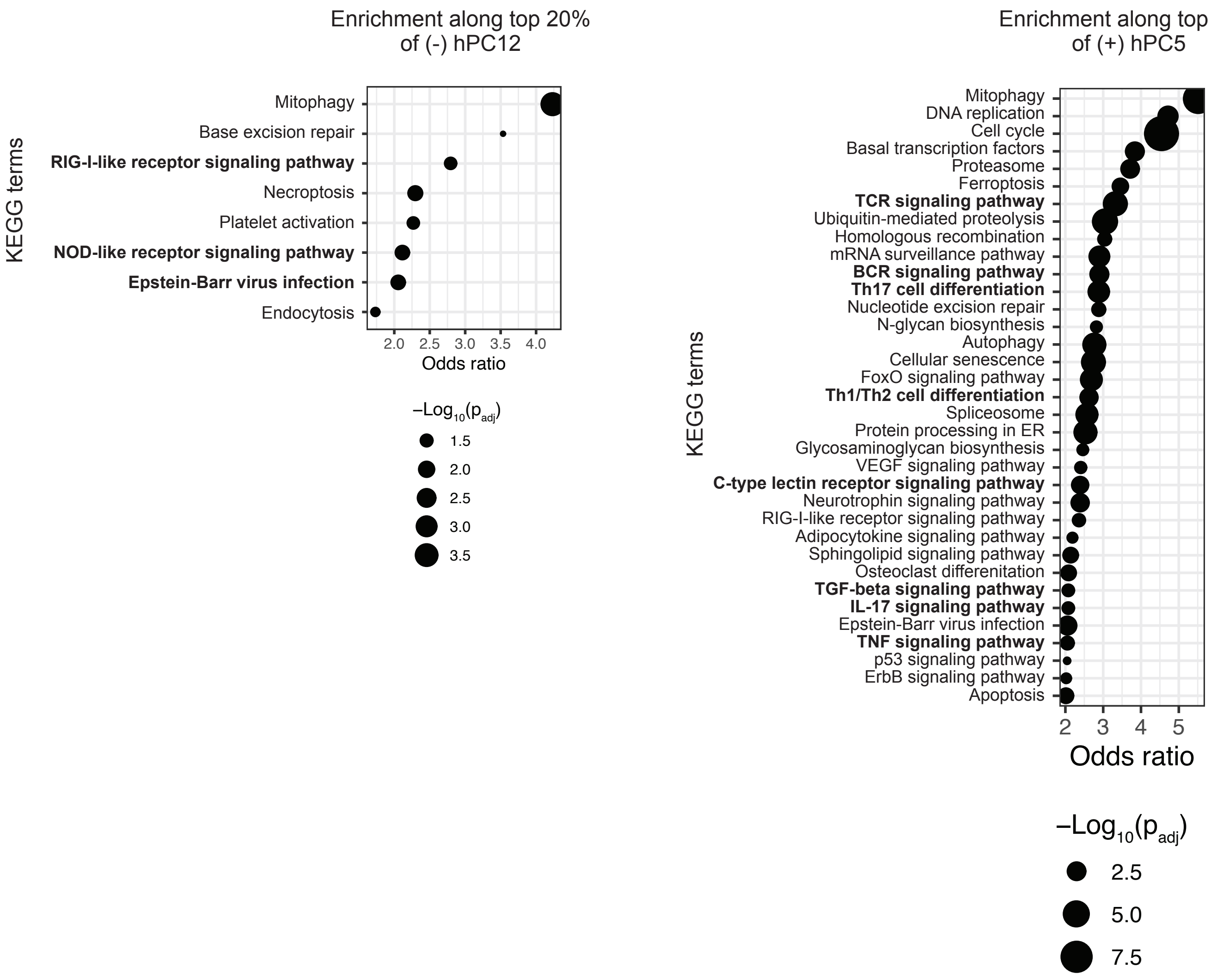


Related to Fig. 4.
KEGG terms significantly enriched on the ‘low CFU’-associated ends of hPC12 (left) and hPC5 (right). Dot size corresponds to adjusted p-value, and terms are plotted according to their odds ratio, which is a statistic that estimates the magnitude of the enrichment.

*Figure S7: ssGSEA on NHP route cohort confirms timepoint-dependent separation of samples by protection status along pathway-associated gene sets of interest*

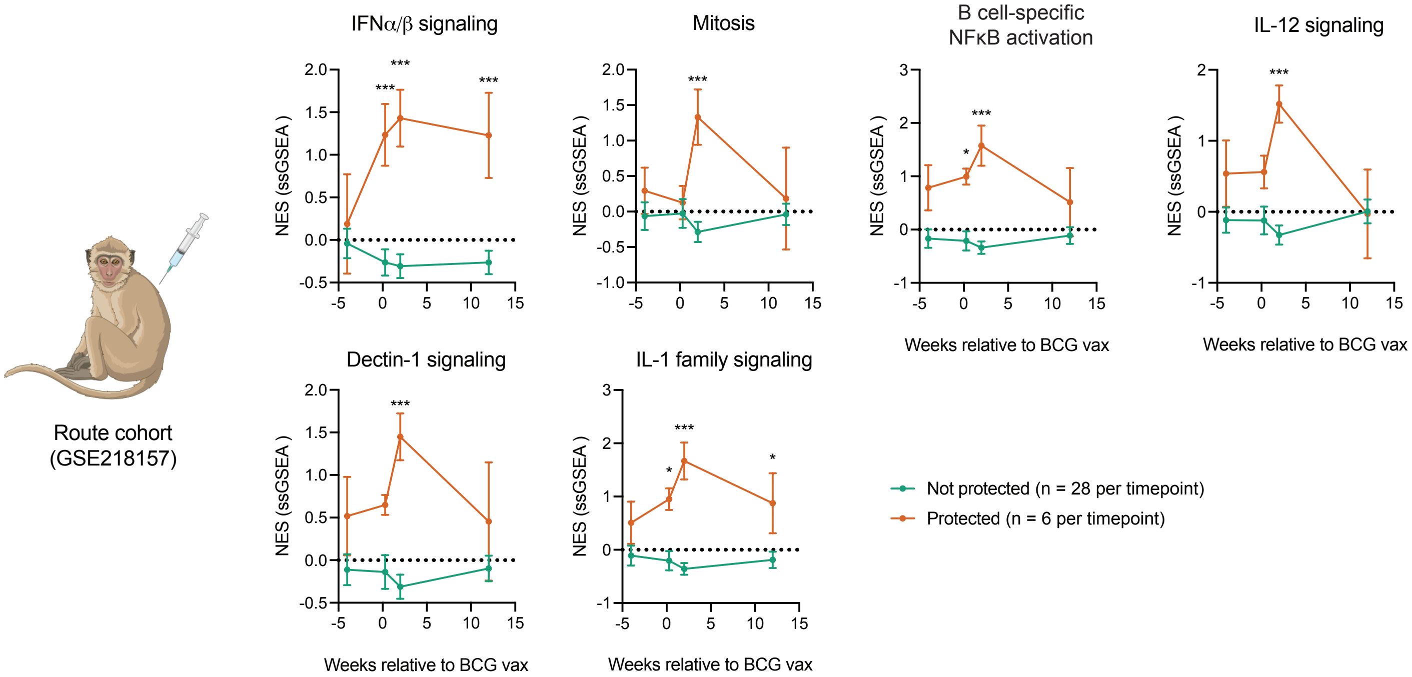


Related to Fig. 4.
Normalized enrichment scores (NES) calculated using single sample gene set enrichment analysis (ssGSEA) for each sample in the NHP route cohort (GSE218157). Gene sets for each pathway/process were derived from the Reactome Pathway Database. Sample scores were then lumped by protection outcome as evaluated post-*Mtb* challenge for visualization. *p < 0.05, **p < 0.01, ***p < 0.001 by unpaired t-test with Holm-Šídák's multiple comparison correction as calculated in GraphPad.
